# NLGC: Network Localized Granger Causality with Application to MEG Directional Functional Connectivity Analysis

**DOI:** 10.1101/2022.03.09.483683

**Authors:** Behrad Soleimani, Proloy Das, I.M. Dushyanthi Karunathilake, Stefanie E. Kuchinsky, Jonathan Z. Simon, Behtash Babadi

**Author notes:** The identification of specific products or scientific instrumentation is considered an integral part of the scientific endeavor and does not constitute endorsement or implied endorsement on the part of the author, DoD, or any component agency. The views expressed in this article are those of the author and do not reflect the official policy of the Department of Army/Navy/Air Force, Department of Defense, or U.S. Government. Email addresses:* (Behrad Soleimani), (Proloy Das), (I.M. Dushyanthi Karunathilake), (Stefanie E. Kuchinsky), (Jonathan Z. Simon), (Behtash Babadi).

## Abstract

Identifying the directed connectivity that underlie networked activity between different cortical areas is critical for understanding the neural mechanisms behind sensory processing. Granger causality (GC) is widely used for this purpose in functional magnetic resonance imaging analysis, but there the temporal resolution is low, making it difficult to capture the millisecond-scale interactions underlying sensory processing. Magnetoencephalography (MEG) has millisecond resolution, but only provides low-dimensional sensor-level linear mixtures of neural sources, which makes GC inference challenging. Conventional methods proceed in two stages: First, cortical sources are estimated from MEG using a source localization technique, followed by GC inference among the estimated sources. However, the spatiotemporal biases in estimating sources propagate into the subsequent GC analysis stage, may result in both false alarms and missing true GC links. Here, we introduce the Network Localized Granger Causality (NLGC) inference paradigm, which models the source dynamics as latent sparse multivariate autoregressive processes and estimates their parameters directly from the MEG measurements, integrated with source localization, and employs the resulting parameter estimates to produce a precise statistical characterization of the detected GC links. We offer several theoretical and algorithmic innovations within NLGC and further examine its utility via comprehensive simulations and application to MEG data from an auditory task involving tone processing from both younger and older participants. Our simulation studies reveal that NLGC is markedly robust with respect to model mismatch, network size, and low signal-to-noise ratio, whereas the conventional two-stage methods result in high false alarms and mis-detections. We also demonstrate the advantages of NLGC in revealing the cortical network-level characterization of neural activity during tone processing and resting state by delineating task- and age-related connectivity changes.

## 1. Introduction

Characterizing the directed connectivity among different cortical areas that underlie brain function is among the key challenges in computational and systems neuroscience, as it plays a key role in revealing the underlying mechanism of cognitive and sensory information processing (Sporns, 2014; Lochmann and Deneve, 2011). A remarkable data-driven methodology for statistical assessment of directed connectivity is commonly referred to as *Granger causality*, which quantifies the flow of information based on improvement in the temporal predictability of a time-series given the history of another one (Bressler and Seth, 2011). Mathematically speaking, for two time series *x*_1,*t*_ and *x*_2,*t*_, if using the history of *x*_1,*t*_ can significantly improve the prediction of *x*_2,*t*_, we say that there is a Granger causal (GC) link from *x*_1,*t*_ to *x*_2,*t*_, i.e., *x*_1_ ↦ *x*_2_,; otherwise, there is no GC link from *x*_1_ to *x*_2_. An essential attribute of Granger causality distinguishing it from other connectivity metrics, such as *Pearson correlation* or *mutual information*, is its directionality, which makes it a powerful statistical tool for brain functional connectivity analysis (Seth et al., 2015).

Granger causality has been widely utilized in analyzing functional magnetic resonance imaging (fMRI) data (Roebroeck et al., 2005; Deshpande et al., 2009; Chen et al., 2018; Dong et al., 2019; Azarmi et al., 2019). In addition to technical challenges such as hemodynamic variability and ambiguity in the interpretation of Granger causality analysis for fMRI data (Roebroeck et al., 2011; Deshpande and Hu, 2012), due to the relatively low temporal resolution of fMRI, on the order of seconds, cortical network interactions that occur on the millisecond-scale in cognitive and sensory processing cannot be captured. Magnetoencephalography (MEG) and Electroencephalography (EEG), on the other hand, provide higher temporal resolution in the order of milliseconds, but unlike fMRI, only provide low-dimensional linear mixtures of the underlying neural sources. Typically, the number of sensors and sources are in the order of ~ 10^2^ and ~ 10^4^, respectively, which makes the problem of estimating cortical sources highly ill-posed (Hämäläinen and Ilmoniemi, 1994; Baillet et al., 2001; Hauk et al., 2019; Samuelsson et al., 2020). To address this issue, existing methods typically follow a two-stage procedure, in which the neuromagnetic inverse problem is solved first to obtain sources estimates, followed by connectivity analysis performed on the estimated sources (Schoffelen and Gross, 2009; Sohrabpour et al., 2016; Brookes et al., 2016; Cope et al., 2017; Farokhzadi et al., 2018; Seymour et al., 2018; Blanco-Elorrieta et al., 2018; Liu et al., 2019, 2020; Rosenberg et al., 2021; Lu et al., 2013; Hejazi and Nasrabadi, 2019; Gao et al., 2020).

While this two-stage approach is convenient to adopt, it comes with significant limitations. First, Granger causality, as a network-level property, is a second-order spatiotemporal relation between two sources. As such, it requires reliable estimates of second-order moments of cortical source activity. Source localization techniques, however, predominantly use strong priors to combat the ill-posedness of the neuromangetic inverse problem and thereby to estimate first-order moments of cortical sources with controlled spatial leakage. In additional to the challenges caused by artefactual spatial mixing and mis-localization of the estimated sources, which can readily complicate connectivity analysis (Palva and Palva, 2012), the biases introduced in favor of accurate estimation of first-order source activities typically propagate to the second stage of connectivity analysis and may result not only in mis-detection of pair-wise interactions, but also capturing spurious ones (Palva et al., 2018).

Second, a necessary step in establishing causal relationships among cortical sources entails accurate estimation of their temporal dependencies. Source localization methods using linear and non-linear state-space models address this challenge by modeling source dynamics as multivariate autoregressive processes (Long et al., 2006; Pirondini et al., 2018; Lamus et al., 2012; Hui and Leahy, 2006; Ding et al., 2007; Limpiti et al., 2009; Nalatore et al., 2009; Sekihara et al., 2010; Cheung et al., 2010; Cheung and Van Veen, 2011; Sekihara et al., 2011; Fukushima et al., 2015; Cho et al., 2015). While these methods are able to notably increase the spatiotemporal resolution of the estimated sources, they come with massive computational requirements, especially when the number of sources and the length of the temporal integration window grows (Long et al., 2011; Cheung et al., 2010; Sekihara et al., 2010). Finally, existing methods that address these challenges lack a precise statistical inference framework to assess the quality of the inferred GC links and control spurious detection (Manomaisaowapak et al., 2021).

In this paper, we address the foregoing challenges by introducing the Network Localized Granger Causality (NLGC) inference framework to directly extract GC links at the cortical source level from MEG data, without requiring an intermediate source localization step. We model the underlying cortical source activity as a latent sparse multivariate vector autoregressive (VAR) process. We then estimate the underlying network parameters via an instance of the Expectation-Maximization (EM) algorithm with favorable computational scalability. The estimated network parameters are then de-biased to correct for biases incurred by the sparsity assumption, and used to form a test statistic that allows to detect GC links with high statistical precision. In doing so, we provide a theoretical analysis of the asymptotic distribution of said test statistic. We evaluate the performance of NLGC through comprehensive simulations by comparing it with several two-stage procedures. Our simulation results indeed confirm the expected performance gains of NLGC in terms of reducing spurious GC link detection and high hit rate.

We further examine the utility of NLGC by application to experimentally recorded MEG data from two conditions of pure-tone listening and resting state in both younger and older individuals. We consider two frequency bands of interest, namely, combined Delta and Theta bands (0.1 – 8 Hz) and Beta band (13 – 25 Hz), for GC analysis which have previously yielded age-related changes in resting state coherence analysis (Fleck et al., 2016). The detected GC networks using NLGC reveal striking differences across the age groups and conditions, in directional interactions between frontal, parietal, and temporal cortices. Further inspection of these networks reveals notable inter-vs. intra-hemispheric connectivity differences. In summary, NLGC can be used as a robust and computationally scalable alternative to existing two-stage connectivity analysis approaches used in MEG analysis.

## 2. Results

### 2.1. Overview of NLGC

Here, we give an overview of the proposed NLGC inference methodology, as depicted in Fig. 1, and highlight the novel contributions.

**Figure 1:**
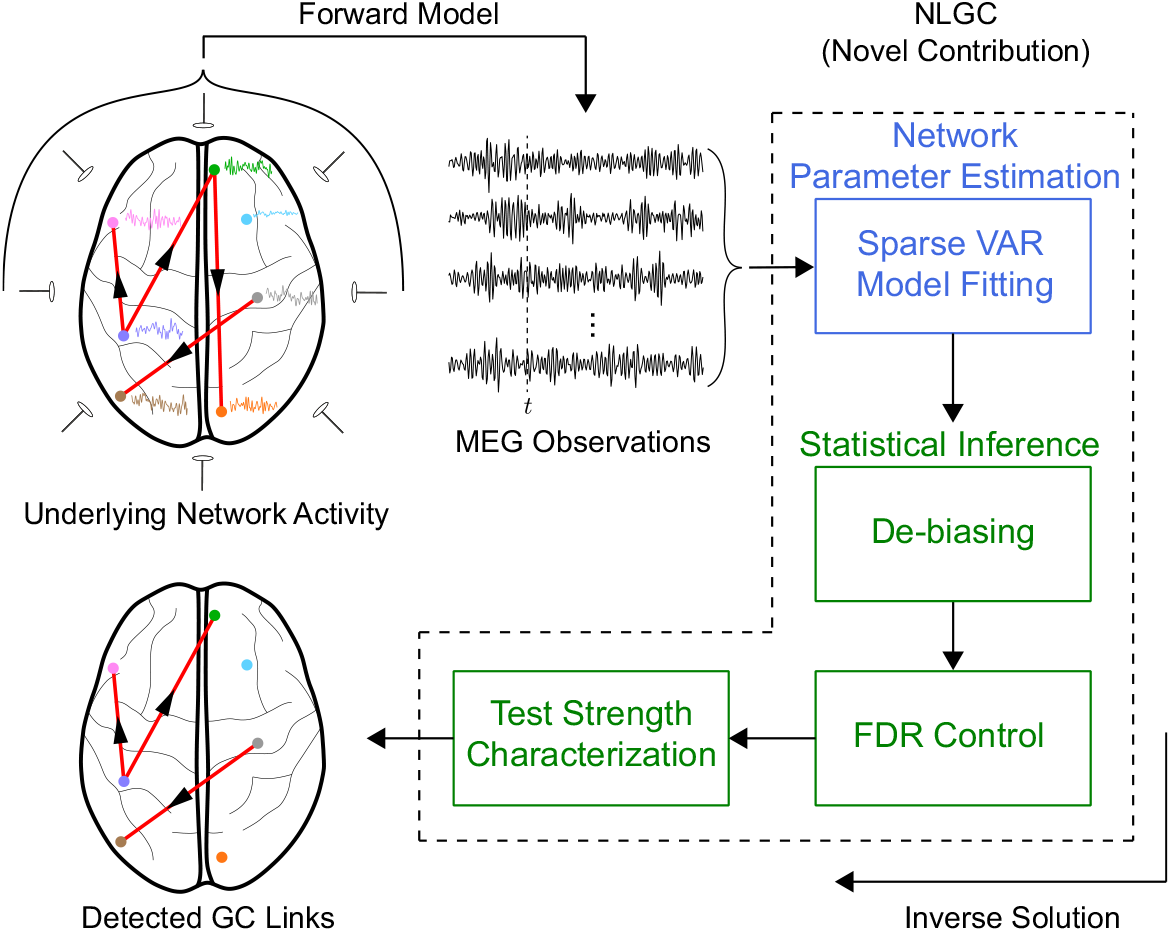
A schematic depiction of the proposed NLGC inference. For cortical sources that form an underlying network, our contribution is to directly infer this network, using the framework of Granger, from the MEG measurements. NLGC is composed of network parameter estimation (blue block) and statistical inference (green blocks) modules. Unlike the conventional two-stage methods, NLGC extracts the GC links without an intermediate source localization step.

The sources of the signals recorded by MEG/EEG sensors are mainly the post-synaptic primary currents of a bundle of tens of thousands of synchronously active pyramidal cells that form an *effective current dipole* (Murakami and Okada, 2006; Hämäläinen et al., 1993; Da Silva, 2009). As such, to formulate the MEG/EEG forward model, a distributed cortical source space is considered in which the cortical surface is discretized using a mesh comprising a finite number of current dipoles placed at its vertices. These current dipoles are henceforth called sources, and their activity as source time-courses.

Assuming that there are *M* such sources, we denote the collective source activity at discrete time *t* as an *M*-dimensional vector **x**_*t*_, where its *i^th^* element, *x_i,t_* is the activity of source *i*, for *i* = 1, 2,⋯, *M* and *t* = 1, 2,⋯, *T*, where *T* denotes the data duration. The *N* MEG sensors measure the *N*-dimensional observation vector **y**_*t*_ at time *t*. The MEG observations follow a well-known linear forward model given by (Sarvas, 1987; Mosher et al., 1999; Baillet et al., 2001):

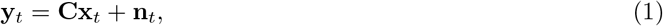

where the *N* × *M* matrix **C** maps the *source space activity* to the *sensor space* and is commonly referred to as the *lead-field matrix.* The *N*-dimensional measurement noise vector **n**_*t*_ is modeled as a zero mean Gaussian random vector with covariance matrix **R** and is assumed to be identically and independently distributed (i.i.d.) across time (Cheung and Van Veen, 2011; Cheung et al., 2010; Long et al., 2011; Wipf et al., 2010).

As for the evolution of the sources, we consider **x**_*t*_ as a latent state vector and model its evolution over time by the following generic stochastic dynamical model:

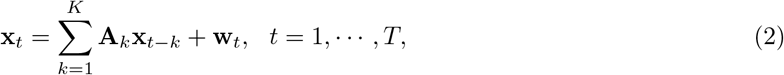

where the *M*-dimensional vectors **w**_*t*_ are assumed to be i.i.d. zero mean Gaussian random vectors with unknown diagonal covariance matrix 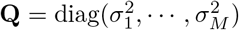 and independent of **v**_*t*_. The *M* × *M* coefficient matrix **A**_*k*_ quantifies the contribution of the neural activity from time *t* – *k* to the current activity at time *t*, for *k* = 1,…, *K*. This dynamical model is conventionally called a Vector Autoregressive (VAR) model of order *K* (or VAR(*K*)) and is commonly used in time-series analysis (Johansen, 1995).

Assuming that the source time-series **x**_*t*_ form an underlying network (Fig. 1, top left), our main contribution is to find the inverse solution to this latent network, in a Granger causality sense, directly from the MEG observations **y**_*t*_ (Fig. 1, bottom left). If reliable estimates of the network parameters 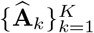 were at hand, one could perform a statistical assessment of causality from source *j* to *i* by checking whether 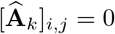 for all *k* =1, 2,⋯, *K* (i.e., no causal link) or 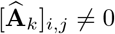 for at least one of *k* = 1, 2,⋯, *K* (i.e., causal link). However, reliable estimation of the network parameters based on noisy and low-dimensional measurements **y**_*t*_ of typically short duration is not straightforward. When noisy, but direct, observations of the sources are available, statistical methods such as LASSO are typically used to test for these hypotheses; however, when the number of sources *M* and lags *K* are large, such methods suffer from the large number of statistical comparisons involved.

The classical notion of Granger causality circumvents this challenge by considering the “bulk” effect of the history of one source on another in terms of temporal predictability. To this end, for testing the GC link from source *j* to source *i*, two competing models are considered: a *full model*, in which all sources are considered in Eq. (2) to estimate the network parameters and thereby predict source *i*; and a *reduced* model, in which the coefficients from source *j* to *i* are removed from Eq. (2), followed by estimating the network parameters and predicting source *i*. The log-ratio of the prediction error variance between the reduced and full models is used as the Granger causality measure. In other words, the better the prediction of the full model compared to the reduced model, the more likely that source *j* has a causal contribution to the activity of source *i*, in the sense of Granger causality.

Considering the inverse problem of Fig. 1, there are several key challenges. First, unlike the classical GC inference frameworks, the sources are not directly observed, but only their low-dimensional and noisy sensor measurements are available. Second, GC inference inherently demands single-trial analysis, but the trial duration of cognitive and sensory experiments are typically short, which renders reliable model parameter estimation difficult. Finally, testing the improvement of the full model over the reduced model requires a precise statistical characterization to limit false detection of GC links.

Existing methods mostly treat these challenges separately, by operating in a two-stage fashion: a source localization procedure is first performed to estimate the sources, followed by performing parameter estimation and conventional GC characterization. However, source localization techniques use specific priors that aim at combating the ill-posed nature of the neuromagnetic inverse problem and thereby bias the source estimates in favor of *spatial* sparsity or smoothness (Lamus et al., 2012; Krishnaswamy et al., 2017; Babadi et al., 2014; Wipf et al., 2010; Sohrabpour et al., 2016; Gramfort et al., 2013b). As such, the network parameters, which inherently depend on second-order current source moments, are recovered from these biased first-order source estimates and thus incur significant errors that complicate downstream statistical analyses.

In contrast, NLGC aims at addressing these challenges jointly and within a unified inference framework. The resulting solution is composed of a network parameter estimation module, in which the VAR model parameters 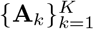 are estimated directly from the MEG data by assuming sparse *interactions* among the sources, as opposed to the commonly-used *spatial* sparsity assumption. As such, the biases induced by this approach only effect the VAR coefficients, and not the spatiotemporal distribution of the sources. Furthermore, we account for these biases in the statistical inference module of NLGC: a de-biasing block is used to correct for biases incurred by sparse VAR estimation, a false discovery rate (FDR) control block is used to correct for multiple comparisons, and a test strength characterization block assigns a summary statistic in the range of [0, 1] to each detected link, denoting the associated statistical test power (i.e., Youden’s *J*-statistic).

While the building blocks that form NLGC are individually well-established in statistical inference literature, including but not limited to Granger causal inference from directly observable states (Bolstad et al., 2011; Endemann et al., 2022) and state-space model parameter estimation (Cheung et al., 2010; Nalatore et al., 2009; Sekihara et al., 2010; Pirondini et al., 2018), our contribution is to unify them within the same framework and specializing them to the problem of direct GC inference from MEG observations. To this end, our technical contributions include: 1) developing a scalable sparse VAR model fitting algorithm by leveraging steady-state approximations to linear Gaussian state-space inference, sparse model selection, and low-rank approximations to the lead field matrix (Sections 4.4.1, 4.5.1, 4.5.2 and Appendix A); and 2) providing a theoretical analysis characterizing the asymptotic distribution of a carefully designed test statistic, namely the de-biased deviance difference, that allows both FDR correction and test strength characterization (Theorem 1 in Section 4.4.3 and Appendix B).

### 2.2. An Illustrative Simulation Study

We first present a simple, yet illustrative, simulated example to showcase how the main components of NLGC work together to address the shortcomings of two-stage approaches. Consider *M* = 84 cortical patches, within which patches 1 through 8 are active and forming a VAR(5) network as shown in Fig. 2A, and the rest are silent (See Section 4.5.1 for details of source space construction). The ground truth GC map of a subset of sources, indexed from 1 through 15, are shown in Fig. 2B (top left) for visual convenience. The (*i,j*) element of the GC matrix indicates the GC link (*j* ↦ *i*). The time courses of the cortical patch activities are observed through a random mixing matrix (each element is independently drawn from a standard normal distribution) corresponding to *N* = 155 sensors for three trials of duration *T* = 1000 samples each. To simulate the MEG observations, we used one lead-field per cortical patch for simplicity. The detailed parameter settings for this simulation study are given in Section 4.8.1.

**Figure 2:**
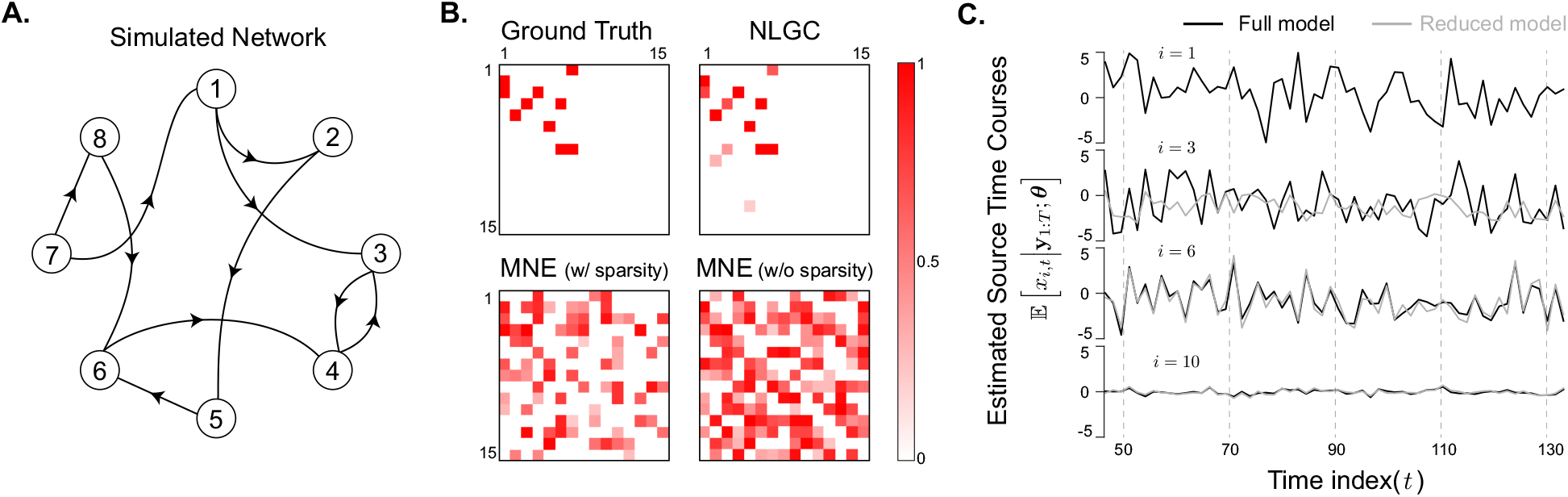
An illustrative simulated example. **A.** The underlying true GC network between the active sources indexed by 1, 2,⋯, 8 (explaining 90% of the power of the 84 sources). The remaining 76 sources are silent and are modeled as independent white noises accounting for the remaining 10% of the source power. **B.** The ground truth and estimated GC maps using NLGC and MNE (with and without accounting for sparsity). Only a subset of sources indexed by 1, 2,⋯, 15 are shown for visual convenience. NLGC fully captures the true links with only a few false detection; on the other hand, the two-stage approaches using MNE, capture around half of the true links, but also detect numerous spurious links. While enforcing sparsity mildly mitigates the false alarm performance of the two-stage approach, it is unable to resolve it. **C**. Estimated activity time-courses of the patches with index 1, 3, 6, and 10 based on full models and the reduced models corresponding to the GC link (1 ↦ 3) and non-GC links (1 ↦ 6) and (1 ↦ 10) as examples. As expected, since the GC link (1 ↦ 3) exists, removing the 1^st^ patch contribution from the VAR model of the 3^rd^ patch dramatically changes the predicted activity of patch 3 (second line). However, this is not the case for the other two examples, since the links (1 ↦ 6) and (1 ↦ 10) do not exist (third and fourth lines).

We compare the performance of NLGC to two baseline two-stage methods composed of an initial source localization stage via the Minimum Norm Estimate (MNE) algorithm, followed by VAR model fitting via either 1) least squares with no sparsity assumption, and 2) *ℓ*_1_-norm regularized least squares to capture sparse parameters, similar to that used in NLGC. The details of the VAR model fitting given the source estimates are presented in Appendix A.3.

Fig. 2B shows the *J*-statistics corresponding to the detected GC links for NLGC and the two baseline methods based on MNE. Note that a *J*-statistic near 1 interprets as a detection with both high sensitivity and specificity, and a *J*-statistic near 0 corresponds to either low sensitivity or specificity, or both. As it can be seen in Fig. 2B, NLGC not only captures the true links, but also only detects a negligible number of false links. On the other hand, the two-stage methods based on MNE only detect about half of the true links and suffer from numerous spurious links. Note that while enforcing sparsity in the two-stage method seems to mitigate the number of spurious links (Fig. 2B, bottom left) compared to the two-stage method with no sparsity (Fig. 2B, bottom right), the errors incurred in the first stage of source localization can not be corrected through the second stage of parameter estimation.

Fig. 2C shows the expected value of estimated cortical patch activities corresponding to the full and reduced models of 4 cortical patches (indexed by 1, 3, 6, and 10). Since the GC link (1 ↦ 3) exists, in the corresponding reduced model, i.e., when the contribution of the 1^st^ cortical patch (shown in the first line) is removed from the VAR model of the 3^rd^ cortical patch, the activity of cortical patch 3 is highly suppressed (second line, gray trace) compared to that of the full model (second line, black trace). On the other hand, for cortical patches 6 and 10, since none of the GC links (1 ↦ 6) and (1 ↦ 10) exist, including or excluding the 1 ^st^ patch in their VAR model does not effect their prediction accuracy and as a result, their estimated activity time-courses for both the full and reduced models are similar (third and fourth lines).

The results so far validate the superior performance of the first component of NLGC, i.e., network parameter estimation. As for the second component, statistical inference, a key theoretical result of this work is to establish the asymptotic distribution of a test statistic called the *de-biased deviance difference* between the full and reduced models of a link (i ↦ *j*), denoted by 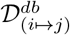 In Theorem 1, we establish that if a GC link from cortical patch *i* to *j* does not exist, the corresponding test statistic 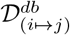 is asymptotically chi-square distributed, and if the GC link exists, 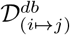 is distributed according to a non-central chi-square.

Here we empirically examine this theoretical result for the foregoing simulation. Consider the links (7 ↦ 1) and (7 ↦ 4) which are GC and non-GC, respectively. We generated 200 different realizations of the VAR processes with the same parameters and compared the empirical distribution of the de-biased deviance corresponding to these two links with their theoretical distribution obtained by Theorem 1. Fig. 3A illustrates the close match between empirical and theoretical distributions of 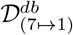 and 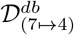 Based on Theorem 1, for the non-GC link (7 ↦ 4), the de-biased deviance has a central χ^2^(5) distribution. On the other hand, the de-biased deviance of the GC link (7 ↦ 1) is distributed according to a non-central χ^2^(5, 61.4).

**Figure 3:**
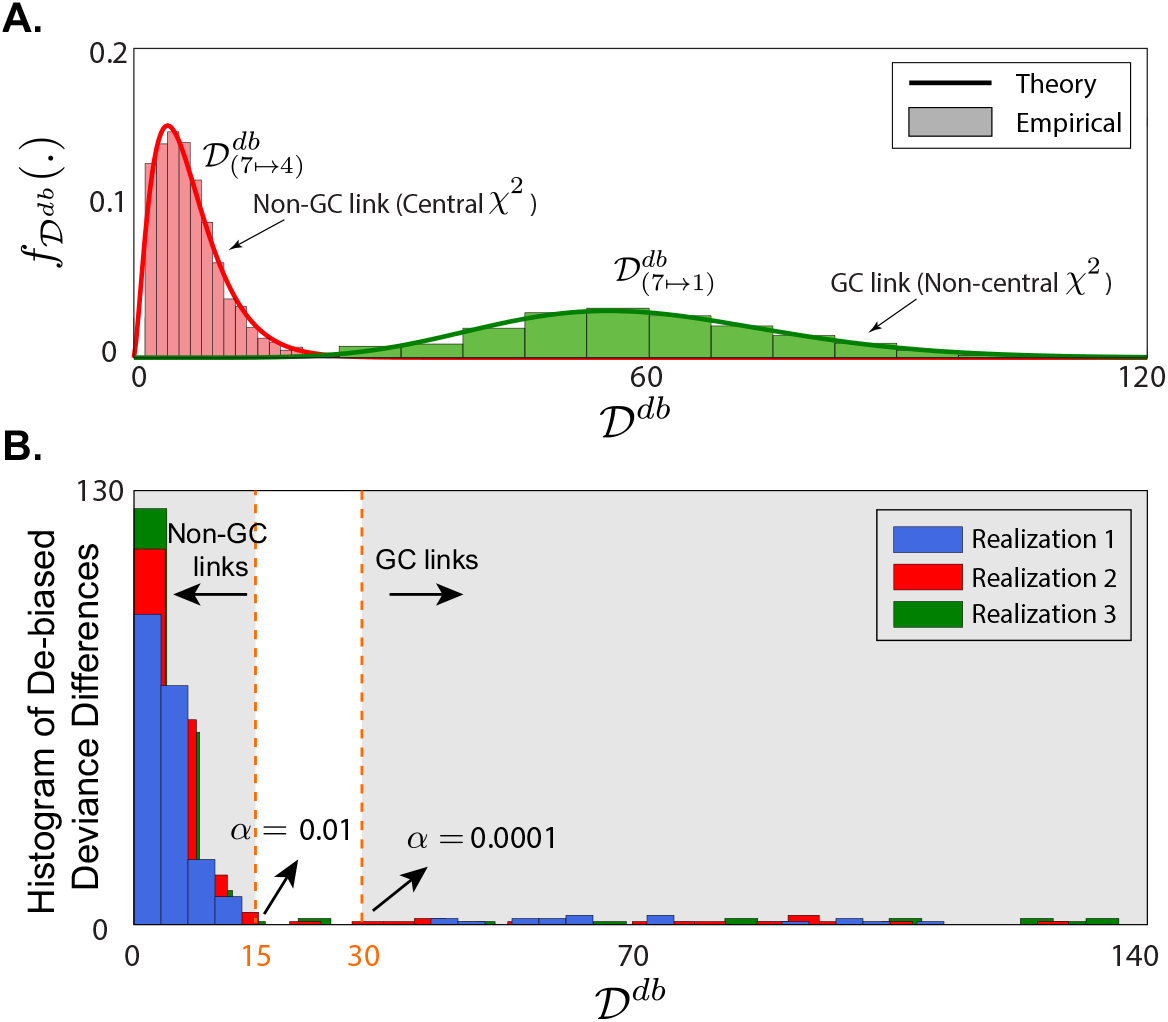
Empirical validation of Theorem 1. **A**. Theoretical and empirical distributions of the de-biased deviance differences corresponding to the GC link (7 ↦ 1) and non-GC link (7 ↦ 4) from the setting of Fig. 2. The empirical distributions closely match the theoretical predictions of Theorem 1. **B.** Histogram of the de-biased deviance differences of all possible links between the first 15 sources for three different realizations of the VAR processes with the same parameters and for two significance levels *α* = 0.01 and 0.0001. The de-biased deviance differences show a clear delineation of the significant GC links (to the right of the dashed vertical lines) and insignificant ones (to the left of the dashed vertical lines), while exhibiting robustness to the choice of the significance level.

In Fig. 3B, the histogram of the de-biased deviance differences corresponding to all links within the subset of sources indexed from 1 through 15 is plotted for three different realizations of the VAR processes with the same parameters as before. Depending on the threshold *α* for rejecting the null hypothesis to detect a GC link, one can obtain an equivalent threshold for 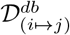. In Fig. 3B, two thresholds are shown with dashed lines for α = 0.01 and 0.0001. It is noteworthy that most of de-biased deviance differences corresponding to the true GC links lie on the right hand side of the dashed lines for both thresholds and for the three realizations, suggesting robustness of GC link detection framework. On the other hand, most of the possible GC links are non-existent in our simulation setting, which results in the concentration of most of the de-biased deviance difference values to the left of the dashed lines, and hence few false detections as shown in Fig. 2B. In NLGC, we further leverage this virtue by using an FDR correction procedure to control the overall false discovery rate at a target level.

### 2.3. Simulated MEG Data Using a Head-Based Model

We next present a more realistic and comprehensive simulation to evaluate the performance of NLGC and compare it with other two-stage approaches based on a number of different source localization techniques. In addition, we consider the effect of signal-to-noise (SNR) ratio and model mismatch on the performance of the different algorithms. The latter is an important evaluation component, as model mismatch is inevitable in practice due to co-registration errors between MR scans and MEG sensors as well as the choice of the distributed cortical source model.

As for the baseline methods, we consider two-stage GC detection schemes in which the source localization is performed by either the classical MNE (Hämäläinen and Ilmoniemi, 1994) and Dynamic Statistical Parametric Mapping (dSPM) (Dale et al., 2000) methods, or the more advanced Champagne algorithm (Wipf et al., 2010). As for the VAR fitting stage, we use the same ℓ_1_-regularized least squares scheme that is utilized by NLGC, to ensure fairness (See Appendix A.3).

In order to create realistic test scenarios for assessing the robustness of the different algorithms, we consider four cases with attributes defined by the presence vs. absence of source model mismatch, and exact vs. relaxed link localization error:

#### Source Model Mismatch

As it is described in detail in Section 4.5.1, in order to reduce the computational complexity of NLGC, we utilize low-rank approximations to the lead field matrix by grouping dipoles over *cortical patches* and summarizing their contribution using singular value decomposition (SVD) to reduce the column-dimension of the lead-field matrix. Let *r*_gen_. be the number of SVD components used for each cortical patch to generate the simulated MEG data, and let *r*_est_. be the number of SVD components used in the GC detection algorithms. Clearly, if *r*_est_. = *r*_gen_., the forward model matches the ones used in the inverse solution, so there is no model mismatch. However, if *r*_est_. < *r*_gen_., some modes of activity in the simulated data cannot be captured by the inverse solution, thus creating a mismatch between the forward and inverse models. We note that this notion of model mismatch pertains to lack of spatial resolution in the inverse model as compared to the forward model. As such, it does not account for the misalignment of the lead-fields with respect to the anatomy, but instead captures the spatial resolution limitation incurred by the choice of the source space used in the inverse solution.

#### Link Localization Error

Suppose that the GC link (*i* ↦ *j*) exists. If in the GC detection algorithm, *i* is mis-localized to *i*′ ≠ *i* or *j* is mis-localized to *j*′ ≠ *j*, the link is considered a miss under the exact link localization error criterion. Let *N*(*k*) be the 6 nearest neighbors of a source *k*. Under the *relaxed* link localization error, if *i*’ ∈ *N*(*i*) and *j*′ ∈ *N*(*j*), we associate (*i*′ ↦ *j*′) to the correct link (*i* ↦ *j*) and consider it a hit. This way, small localization errors, potentially due to errors in the head model or the underlying algorithms can be tolerated.

The source space is again composed of *M* = 84 cortical patches whose activity is mapped to *N* = 155 MEG sensors using a real head model from one of the subjects in the study. For more details on the parameter settings for this study, see Section 4.8.2. Fig. 4A shows the ground truth GC network and the estimated ones using NLGC and two-stage methods using MNE, dSPM, and Champagne when *m* = 10 patches are active. In this case, NLGC detected no spurious links and missed only 3 of the true GC links. On the other hand, even though MNE, dSPM and Champagne capture almost all true GC links, they suffer from a considerable number of falsely detected GC links.

**Figure 4:**
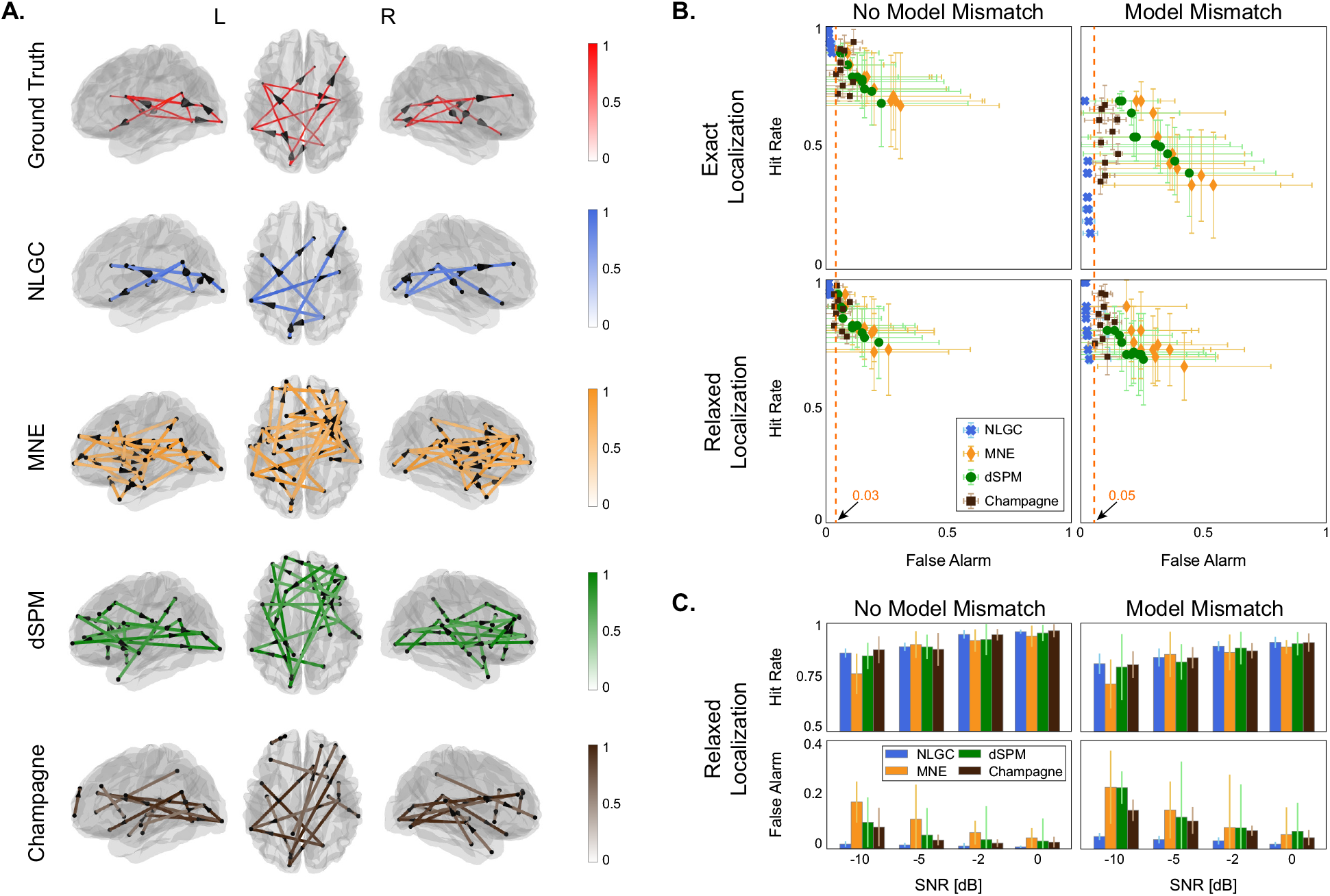
Comparison of NLGC with two-stage procedures using a realistic simulation setting. **A.** Example of the ground truth GC network, and estimates obtained by NLGC and two-stage approaches based on MNE, dSPM, and Champagne overlaid on dorsal and lateral brain plots, with *m* = 10 active patches. NLGC captures nearly all the existing GC links with no spurious detection, whereas the other three methods suffer from significant false detection. **B.** ROC curves (hit rate vs. false alarm) corresponding to NLGC, and two-stage approaches based on MNE, dSPM, and Champagne for exact/relaxed link localization and in the presence/absence of model mismatch. Each point corresponds to simulating data based on m active patches averaged over 10 different realization with randomly assigned source locations, for *m* = 2, 4,⋯, 20. NLGC provides equal or better hit rate, while consistently maintaining low false alarm rate. **C.** Evaluating the effect of SNR for an example setting of *m* = 12 active patches in presence/absence of model mismatch. While the hit rate of NLGC is comparable or better than the other algorithms, it consistently maintains low false alarm rates across a wide range of SNR settings.

To quantify this further, Fig. 4B shows the receiver operating characteristic (ROC) curves corresponding to the different methods for exact vs. relaxed link localization and presence vs. absence of model mismatch. Each point is obtained by varying the number of active patches *m* in the simulation in the range *m* = 2,4,⋯, 20 and averaging the performance of each method over 10 independent trials with randomly allocated patch locations. The 95% quantiles for the hit and false alarm rates are shown as vertical and horizontal bars, respectively. In the absence of source model mismatch (left columns), NLGC outperforms the other three methods in terms of both hit and false alarm rates. The gap between NLGC and the other methods widens when there is source model mismatch (right column, top panel). While the hit rate of NLGC degrades using the exact localization criterion, it remarkably maintains a false alarm rate of < 5%, whereas the other algorithms exhibit false alarm rates as high as ~ 50%. By using the relaxed link localization error criterion (bottom plots), the hit rate of NLGC becomes comparable or better than the other three methods, while it still maintains its negligible false alarm rate. Moreover, the corresponding vertical and horizontal errors bars for NLGC are considerably smaller than the other three algorithms, suggesting the robustness of NLGC to the location of the active patches used for different trials.

Finally, in Fig. 4C, the hit and false alarm rates are plotted for varying levels of SNR in the range {0, −2, −5, −10} dB. The performance is averaged over 10 trials for *m* =12 active patches. As the SNR reduces, even though the performance of all four methods becomes similar in terms of the hit rate, NLGC maintains its low false alarm rate whereas the other algorithms exhibit considerably high rates of false alarm.

Overall, while NLGC achieves comparable hit rate to the other three methods, it maintains consistently low false alarm rates over a wide range of the simulation parameter space. This is a highly desirable virtue, as false detection is the main pitfall of any connectivity analysis methodology. Thus, this simulation study corroborates our assertion that NLGC is a reliable alternative to existing two-stage approaches.

### 2.4. Application to Experimentally Recorded MEG Data

We next consider application to MEG data from auditory experiments involving both younger and older subjects (the data used here is part of a larger experiment whose results will be reported separately). The MEG data corresponds to recordings from 22 subjects, 13 younger adults (5 males; mean age 21.1 years, range 17–26 years) and 9 older adults (3 males; mean age 69.6 years, range 66–78 years). Resting state data were recorded before and after the main auditory task, each 90 s long in duration. During the resting state condition, subjects with eyes open fixated at a red cross at the center of a grey screen. Just before the first resting state recording, 100 repetitions of 500 Hz tone pips were presented, during which the subjects fixated on a cartoon face image at the center of the screen and were asked to silently count the number of tone pips. The tones were presented at a duration of 400 ms with a variable interstimulus interval (1400, 1200, and 1000 ms). The task was around 150 s long, from which two segments, each 40 s long in duration, were used for analysis. More details on the experimental setting is given in Section 4.6.

In order to assess the underlying cortical networks involved in tone processing and compare them with the resting sate, we further considered two key frequency bands of interest (Shafiei et al., 2021), namely the combined Delta and Theta bands (0.1–8 Hz), here called Delta+Theta band, and the Beta band (13– 25 Hz). Since the goal is to capture the (age-related) differences across tone listening versus resting state conditions, we combined the Delta and Theta bands for simplicity of our analysis, as they are both shown to be primarily involved in auditory processing (Baar et al., 2001). In addition, to structure our analysis in an interpretable fashion, we considered the frontal, temporal, and parietal regions of interest (ROIs) in each hemisphere, which are known to play key roles in auditory processing and to change with age (Kuchinsky and Vaden, 2020).

#### *NLGC for the Delta+Theta Band (*0.1 – 8 *Hz)*

Fig. 5A shows the detected GC links between frontal (F) and temporal (T) areas overlaid on the dorsal brain view, for the tone processing vs. resting state conditions and separately for the younger and older subjects. The group average of the detected links across younger and older participants are shown on the left and those of two representative individuals (one younger and one older) are shown on the right. Note that the links involving parietal areas are not shown for the sake of visual convenience. As it can be seen from both the group average and individual-level plots, the top-down links from frontal to temporal areas (red arrows) have a higher contribution to tone processing (first and third columns) compared to resting state (second and fourth columns) for both younger and older adults. On the other hand, more bottom-up links from temporal to frontal areas (green arrows) are detected in the resting state as compared to the tone processing condition.

**Figure 5:**
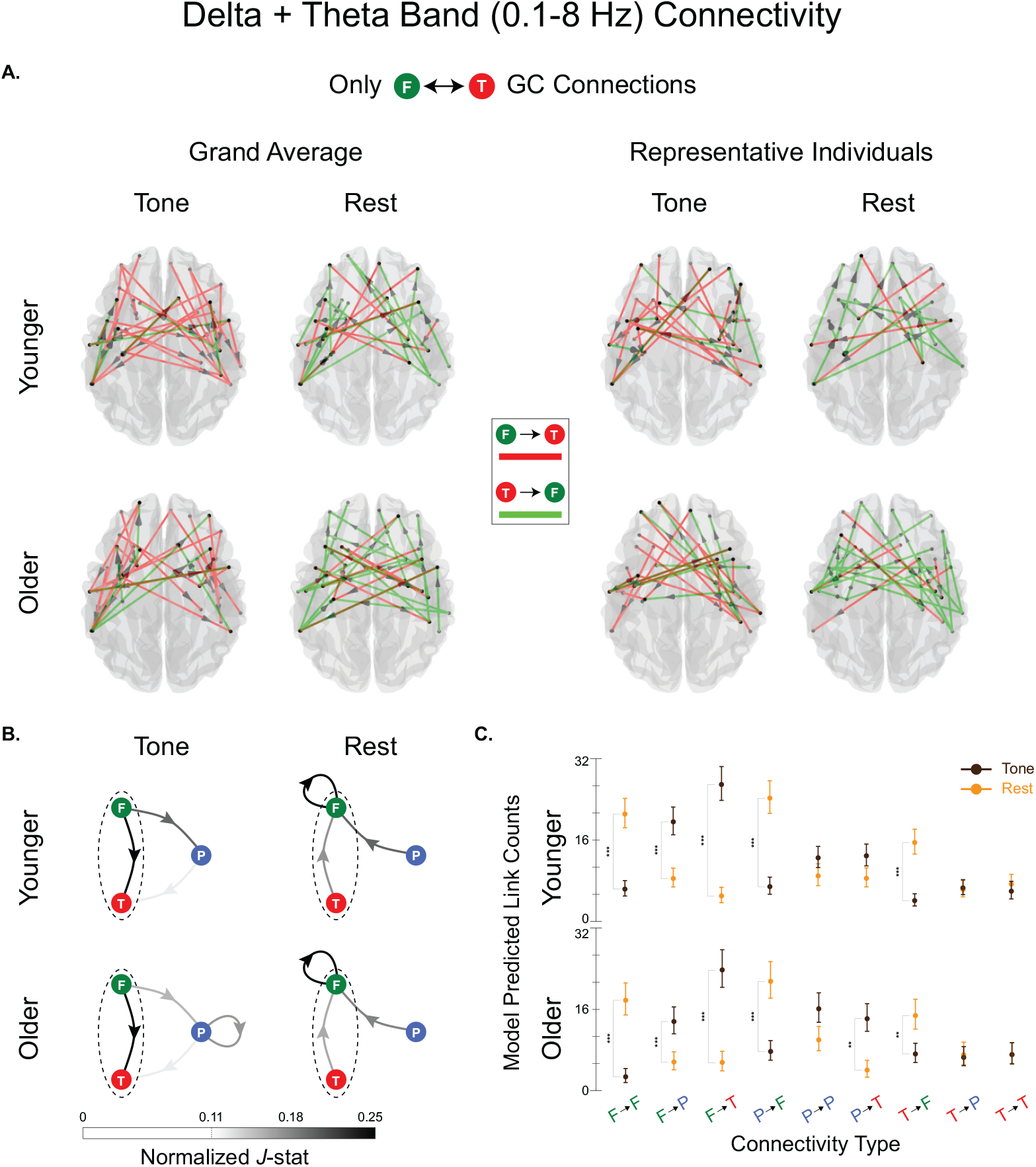
NLGC analysis of experimentally recorded MEG data in the Delta+Theta band (0.1 – 8 Hz). **A.** Extracted GC links between frontal and temporal areas overlaid on dorsal brain plots for younger (top row) and older (bottom row) participants. The first two columns correspond to the group averages and the last two correspond to two representative participants, for the two task conditions of tone processing (first and third columns) and resting state (second and fourth columns). For the group average plots, only *J*-statistic values greater than 0.75 are shown for visual convenience. There is a notable increase of top-down links from frontal to temporal areas during tone processing (red arrows, first and third columns) as compared to the resting state in which bottom-up links from temporal to frontal areas dominate (green arrows, second and fourth columns). **B.** Normalized *J*-statistics, averaged over subjects within each age group, between frontal, temporal, and parietal areas for tone processing vs. resting state conditions and younger vs. older participants. The dashed ovals indicate the normalized average number of links shown in panel A. There are notable changes across task conditions, including dominantly top-down frontal to temporal/parietal connections during tone processing, in contrast to dominantly bottom-up temporal/parietal to frontal connections during resting state. **C.** Statistical testing results showing several significant differences across conditions. No significant age difference is detected in the Delta+Theta band (\*\*\**p* < 0.001; \*\**p* < 0.01; \**p* < 0.05).

In Fig. 5B, the average normalized *J*-statistics of the detected GC links between the frontal, temporal and parietal (P) ROIs are shown as color-weighted edges in a directed graph. For instance, the arrows between temporal and frontal areas, enclosed in dashed ovals, show the normalized average of the arrows shown in the first two columns of Fig. 5A. In addition to the notable change of connectivity between temporal and frontal areas, i.e., from dominantly bottom-up under resting state to dominantly top-down under tone processing, there are several other striking changes both across conditions and age groups. First, from tone processing to the resting state condition, for both age groups, the contribution of outgoing links from frontal to parietal and temporal areas drops. Secondly, in the resting state condition, incoming GC links from parietal and temporal to frontal areas increase. Finally, frontal to frontal interactions become more prevalent in the resting state condition, for both younger and older subjects.

To further quantify these observation, Fig. 5C summarizes statistical test results for comparing the detected link counts for the different connectivity types and across age groups. Interestingly, no significant difference between younger and older participants is detected in either of the conditions. Within each age group, however, several significant changes are detected. In particular, the aforementioned visual observations from Fig. 5B are indeed statistically significant: the top-down frontal to temporal connectivity under tone processing switches to bottom-up temporal to frontal connectivity; outgoing links from the frontal to temporal/parietal areas are significantly increased under tone listening compared to resting state; parietal to frontal connections have more contribution in the resting state compared to tone processing; and frontal to frontal connections increase in the resting state, as previously reported in the literature (Müller et al., 2009; Di Liberto et al., 2018; Henry et al., 2017).

We further inspected the inter- vs. intra-hemispheric contributions of the aforementioned changes, as shown in Fig. 6, where we have combined the older and younger subject pools, given that no significant age difference was detected. In the resting state, the inter- and intra-hemispheric networks are similar (Fig. 6A, right column). However, there are several interesting changes in the inter- vs. intra-hemispheric networks under tone processing (Fig. 6A, left column), such as the increased involvement of intra-hemispheric connections from frontal to parietal and from parietal to temporal areas. Statistical test results shown in Fig. 6B suggest that the detected intra-hemispheric connections are significantly higher than inter-hemispheric ones under tone processing. In addition, the change from a dominantly bottom-up temporal to frontal network under resting state to a dominantly top-down frontal to temporal network under tone processing occurs at both inter- and intra-hemispheric levels.

**Figure 6:**
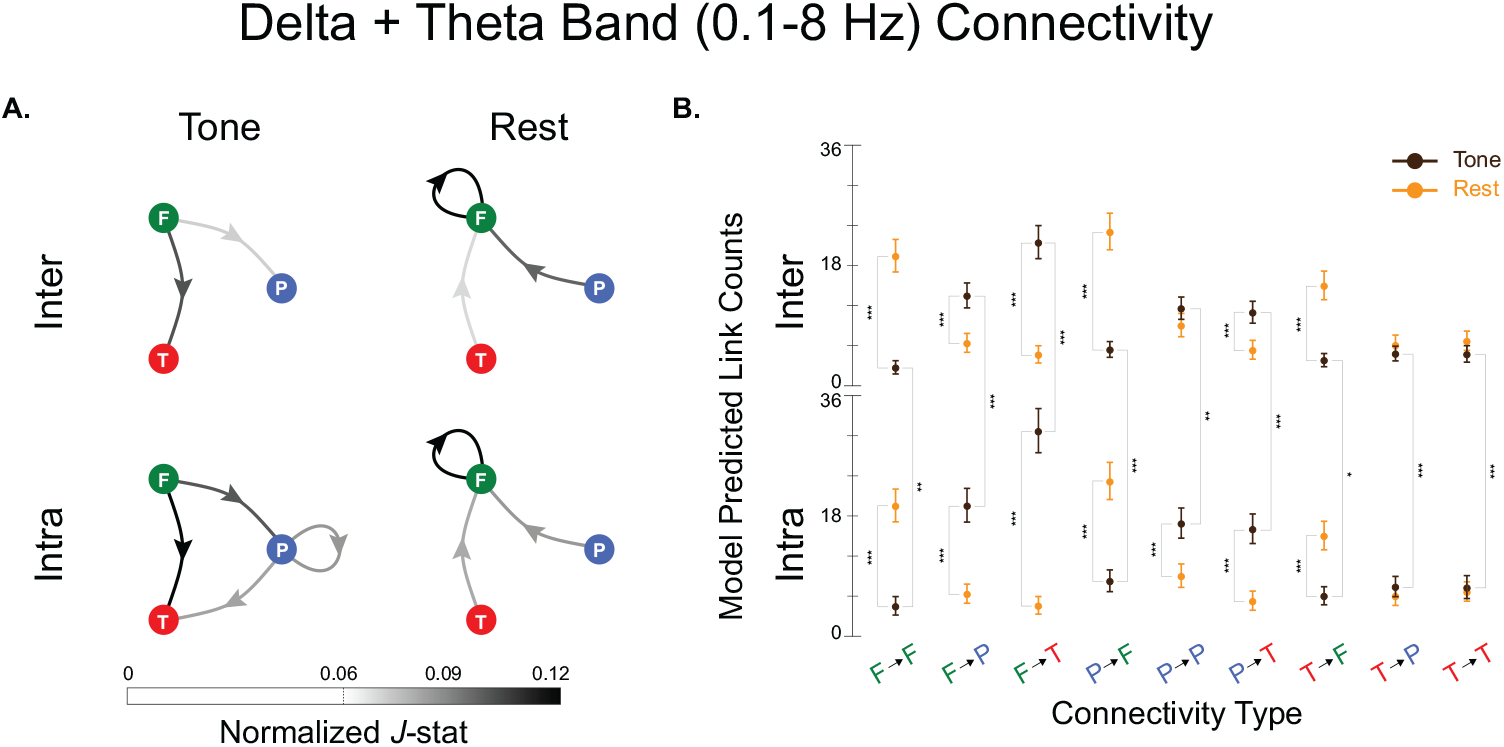
Inter- vs. intra-hemispheric refinement of the analysis of experimentally recorded MEG data in the Delta+Theta band (0.1 – 8 Hz). **A.** Normalized *J*-statistics, averaged over all subject, between frontal, temporal, and parietal areas for inter-hemispheric and intra-hemispheric connectivity types. Given that no significant age difference was detected, the two age groups are pooled together. While the inter- vs. intra-hemispheric contributions to the detect networks are highly similar under resting state, there notable differences under tone processing, including higher number of intra-hemispheric connections from frontal to parietal and from parietal to temporal areas. **C.** Statistical testing results showing several significant differences across conditions and inter- vs. intra-hemispheric contributions (\*\*\**p* < 0.001; \*\**p* < 0.01; \**p* < 0.05).

#### *NLGC for the Beta Band (*13 – 25 *Hz)*

Fig. 7 shows the results of Beta band NLGC analysis in a similar layout as Fig. 5. Fig. 7A shows the detected GC links between frontal and parietal areas for the tone processing vs. resting state conditions and separately for the younger and older subjects. The group average of the detected links across younger and older participants are shown on the left and those of two representative individuals (one younger and one older) are shown on the right. Note that the links involving temporal areas are not shown for the sake of visual convenience. As it can be seen from both the group average and individual-level plots, there is a striking dominance of frontal to parietal links (blue arrows) for older subject under tone listening (first and third columns, bottom plots), whereas in all the other three cases, parietal to frontal links (green arrows) dominate.

**Figure 7:**
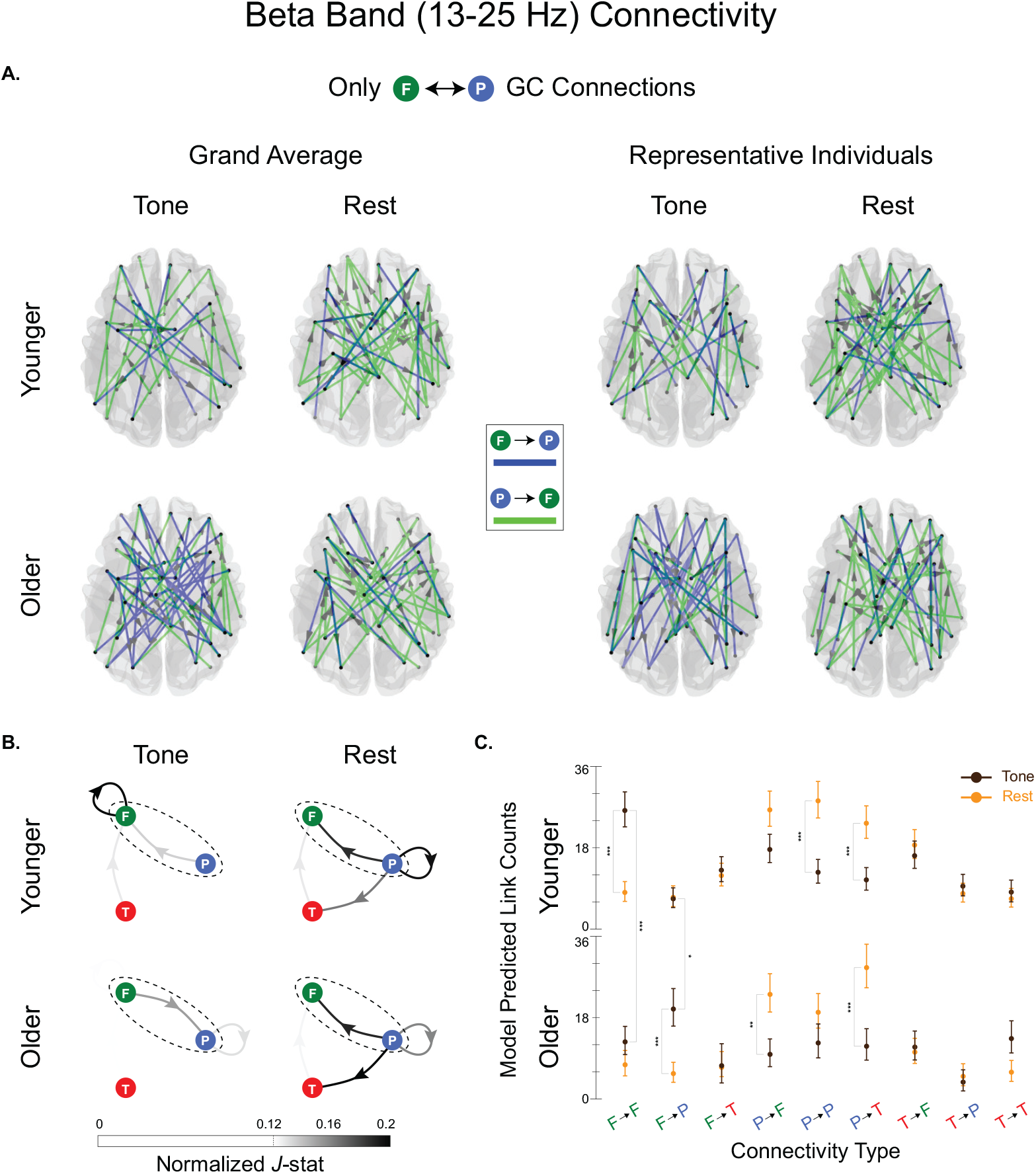
NLGC analysis of experimentally recorded MEG data in the Beta band (13 – 25 Hz). **A.** Extracted GC links between frontal and parietal areas overlaid on dorsal brain plots for younger (top row) and older (bottom row) participants. The first two columns correspond to the group averages and the last two correspond to two representative participants, for the two task conditions of tone processing (first and third columns) and resting state (second and fourth columns). For the group average plots, only *J*-statistic values greater than 0.75 are shown for visual convenience. There is a notable increase of frontal to parietal links under tone processing for older adults (blue arrows, first and third columns, bottom row), whereas in all the other cases parietal to frontal links (green arrows) are dominant. **B.** Normalized *J*-statistics, averaged over subjects within each age group, between frontal, temporal, and parietal areas for tone processing vs. resting state conditions and younger vs. older participants. The dashed ovals indicate the normalized average number of links shown in panel A. There are notable changes across both task conditions and age groups, including the higher involvement of parietal areas during resting state, increase of frontal to frontal connections for younger participants and top-down links from frontal to parietal areas for older participants, during tone processing. **C.** Statistical testing results showing several significant differences across task conditions and age groups (\*\*\**p* < 0.001; \*\**p* < 0.01; \**p* < 0.05).

Fig. 7B shows the average normalized *J*-statistics of the detected GC links between the frontal, temporal and parietal ROIs as color-weighted edges in a directed graph. The edges between parietal and frontal areas, enclosed in dashed ovals, correspond to the normalized average of the weighted arrows shown in the first two columns of Fig. 7A. The GC network under the resting state condition is similar for both age groups, but during tone processing, the network structures are quite different. First, for younger subjects, frontal to frontal connections have a higher contribution to the network as compared to older subjects. On the other hand, as pointed out earlier, for older participants during tone processing, the number of incoming links to parietal from frontal areas increase, as compared to the younger group. Finally, for both younger and older subjects, there are more parietal to temporal connections in resting state compared to tone processing. Fig. 7C summarizes the statistical test results which indeed show both across-age and across-condition differences, for the two connectivity types of frontal to frontal and frontal to parietal, as well as several connectivity changes across the task conditions within the two age groups.

## 3. Discussion and Concluding Remarks

Extracting causal influences across cortical areas in the brain from neuroimaging data is key to revealing the flow of information during cognitive and sensory processing. While techniques such as EEG and MEG offer temporal resolution in the order of milliseconds and are thus well-suited to capture these processes at high temporal resolution, they only provide low-dimensional and noisy mixtures of neural activity. The common approach for assessing cortical connectivity proceeds in two stages: first the neuromagnetic inverse problem is solved to estimate the source activity, followed by performing connectivity analysis using these source estimates. While convenient to use, this methodology suffers from the destructive propagation of the biases that are introduced in favor of source localization in the first stage to the second stage of network inference, often resulting in significant spurious detection.

In this work, we propose a unified framework, NLGC inference, to directly capture Granger causal links between cortical sources from MEG measurements, without the need for an intermediate source localization stage and with high statistical precision. We evaluated the performance of NLGC through comprehensive simulation studies, which revealed the performance gains of NLGC compared to the conventional two-stage procedures in terms of achieving high hit rate, remarkably low false alarm rate, and robustness to model mismatch and low SNR conditions.

We applied NLGC to experimentally recorded MEG data from an auditory experiment comparing trials of tone processing and resting conditions, from both younger and older participants. We analyzed the data in two frequency bands whose coherence has been shown to differ when processing auditory stimuli compared to rest (Weiss and Rappelsberger, 2000), namely the combined Delta+Theta band and the Beta band. The extracted cortical networks using NLGC revealed several striking differences across the frequency bands, age groups, and task conditions. In particular, in the Delta+Theta band, the networks were dominantly top-down from frontal to temporal and parietal areas during tone processing. Previous studies have observed increased coherence between frontal and central and temporal electrodes during auditory processing versus rest, potentially indicative of greater demands on memory and inhibitory processes that are required for active listening (Weiss and Rappelsberger, 2000). Greater anterior to posterior interactivity has particularly been observed in the Theta band in support of working memory (Sarnthein et al., 1998) and other top-down processes (Sauseng et al., 2008), in line with the functioning of the frontal-parietal attention network (Sauseng et al., 2005). However, during resting state, bottom-up links towards frontal areas significantly increased. This broadly aligns with a previous Granger causality analysis that found evidence of unidirectional parietal to frontal connections during resting state fMRI (Duggento et al., 2018). In addition, intra-hemispheric links were more dominant during tone processing as compared to inter-hemishpheric links, whereas the inter- and intra-hemispheric contributions were nearly balanced during resting state. This may align with evidence that even low level auditory stimuli are processed in a lateralized fashion (Millen et al., 1995; Brown and Nicholls, 1997). Additionally, in an fMRI study of 100 adults, Granger causality analyses revealed that parietal-to-frontal connectivity was localized to within-hemispheric pathways (Duggento et al., 2018). Cross-hemispheric connectivity was largely observed within lobes (e.g., frontal-to-frontal). Although there are a number of methodological differences between these studies, together they suggest that NLGC can reveal robust differences in the directionality and band specificity of patterns of connectivity during task processing and at rest.

In general, greater and/or more extensive frontotemporalparietal functional connectivity has been observed when processing clearer auditory stimuli (Abrams et al., 2013; Yue et al., 2013) and for younger compared to older adults (Andrews-Hanna et al., 2007; Peelle et al., 2010). The current results broadly align with these results, but further indicate the directionality and frequency band that may drive those observed differences in connectivity. While our analysis of the Delta+Theta band did not suggest any age differences across age groups, the networks seen in the Beta band revealed key age-related differences during the tone processing task. For younger participants, most of the connections were from parietal and temporal to frontal areas, including frontal to frontal connectivity. However, in older participants, parietal areas were significantly more engaged in the network with notable connections towards frontal areas. Long-range synchrony between frontal and parietal cortices in the Beta band has been observed to dominate during top-down attentional processing (Buschman and Miller, 2007) and is thought to support the enhancement of task-relevant information (Antzoulatos and Miller, 2016). There is also some evidence that Beta band connectivity increases with aging (Moezzi et al., 2019; Vysata et al., 2014). The results did not yield support for previous observations of inter-hemispheric asymmetry reduction with age (Dolcos et al., 2002) in terms of increasing inter-hemispheric connectivity (Maurits et al., 2006). However, this is likely due to the simplicity of the tone counting and rest conditions examined in the present study. Future analyses of speech materials with greater task demands may be more sensitive to such differences.

The NLGC framework includes several technical contributions that are unified within the same methodology, but may also be of independent interest in neural signal processing. These include: 1) a scalable sparse VAR model fitting algorithm based on indirect and low-dimensional observations, that leverages steady-state approximations to linear Gaussian state-space inference, sparse model selection, and low-rank approximations to the lead field matrix; and 2) establishing the asymptotic distributions of the de-biased deviance difference statistics from MEG observations, that may be used in more general hypothesis testing frameworks.

Along with its several improvements over existing work, NLGC comes with its own limitations. First, NLGC requires sufficiently long trial duration, so that the underlying network parameters can be estimated reliably. While the sparsity regularization in NLGC mitigates this issue to some extent, in general the number of parameters needed to be estimated from *NT* observed MEG sensor data points is in the order of ~ *K M*^2^. As an example, to ensure that the number of parameters is in the order of the number of data points for the sake of estimation accuracy, for the typical configurations in this work (i.e., *N* = 155 sensors, *M* = 84 sources, 5-fold cross-validation, 10 Hz frequency band, 100 ms integration window), trials of at least *T* = 25 s in duration are needed. While this requirement was satisfied by the experimental trials used in our work, as also validated in Section 4.8.3, NLGC may not perform well in experiments involving short trials, such as those studying sensory evoked field potentials in which a large number of trials, each in the order of 1 s in duration, are available (David et al., 2006a,b).

Second, while NLGC maintains a remarkably low false alarm rate in a wide range of settings, it is more sensitive to model mismatch in terms of its hit rate performance, as compared with existing two-stage approaches, as examined in Fig. 4B. This is due to the fact that while integrating source localization and VAR parameter estimation in NLGC is advantageous to rejecting spurious GC links, eliminating the first stage of source localization makes NLGC more sensitive to the accuracy of the source space used in estimating the source time-courses and thereby correctly detecting the true GC links. The hit rate performance of NLGC could be improved by using a more refined source space, but this in turn might require a longer observation duration for accurate parameter estimation. Finally, our experimental data validation here was limited by the lack of access to ground truth source activity. We defer validating the performance of NLGC using invasive recordings such as electrocorticography or intracranial EEG, in which the sources are directly observable, to future work.

In addition to the aforementioned technical contributions, NLGC also offers several practical advantages over existing work. First, due to its scalable design, it can be applied to any frequency band of interest to extract the underlying GC networks. Secondly, due to the precise statistical characterization of the detected links, the networks can be transformed to span ROIs of arbitrary spatial resolution, from cortical dipoles to anatomical ROIs, cortical lobes, and hemispheres. Third, unlike most existing connectivity analysis methods that require heavy trial averaging to mitigate spurious detection, NLGC exhibits robustness to model mismatch and low SNR conditions, even where few trials are available. Finally, thanks to the plug-and-play nature of the NLGC building blocks, it can be modified for inferring other network-level characterizations, such as cortical transfer entropy (Daube et al., 2022). To ease reproducibility, we have made a python implementation of NLGC publicly available on Github (Soleimani and Das, 2022). In summary, NLGC can be used as a robust and scalable alternative to existing approaches for GC inference from neuroimaging data.

## 4. Theory and Methods

Here we lay out in detail the generative framework that entails the computational model for relating the neural activity, which produces magnetic fields outside of the brain, to the recordings at the highly sensitive MEG sensors. This generative framework deals with the unobserved neural activity as latent entities: the notion of Granger causality is defined with respect to the latent neural activity. We then propose a novel approach to identify the parameters of the generative model from the multi-channel MEG recordings and construct Granger causal measures to quantify the detected links. We call this unified framework the Network Localized Granger Causality (NLGC) framework.

### 4.1. Main Problem Formulation

Recall the observation and state evolution models given in Eqs. (1) and (2):

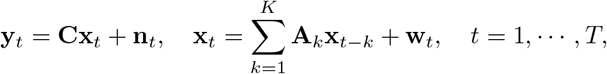

where *T* is the observation duration, **x**_*t*_ ∈ ℝ^*M*^ and **y**_*t*_ ∈ ℝ^*N*^ are, respectively, the cortical activity of *M* distributed sources and the measurements of *N* sensors at time *t*. The process noise **w**_*t*_ and observation noise **v**_*t*_ are assumed to be independent of each other and are modeled as i.i.d. sequences of zero mean Gaussian random vectors with respective covariance matrices 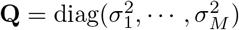 and **R**.

The lead-field matrix **C** ∈ ℝ^*N*×*M*^ can be estimated using a quasi-static solution to the Maxwell’s equations using a realistic head model obtained by MR scans (Sarvas, 1987; Mosher et al., 1999; Baillet et al., 2001). The measurement noise covariance matrix **R** is assumed to be known, as it can be estimated based on empty room recordings (Engemann and Gramfort, 2015). Thus the unknown parameters in these models are: the *M* × *M* coefficient matrices **A**_*k*_, that quantify the contribution of the neural activity from time *t* – *k* to the current activity at time *t*, for *k* = 1,…, *K*, and the process noise covariance matrix **Q**.

Assuming that the source time-series **x**_*t*_ form an underlying network, our main contribution is to find the inverse solution to this latent network, in the sense of Granger causality, directly from the MEG observations **y**_*t*_. We first give an overview of Granger causality while highlighting the challenges in GC inference from MEG data.

### 4.2. Overview of Granger Causality

First, we assume that the sources **x**_*t*_ are directly observable. Noting that [**A**_*k*_]_*i,j*_ quantifies the contribution of source *j* at time *t* – *k* to the present activity of source *i* at time *t*, one can statistically assess the causal effect of source *j* on source *i* via the following hypothesis test:

- *H*_0_: [**A**_*k*_]_*i,j*_ =0 for all *k* = 1, 2,⋯, *K*, i.e., there is no causal influence from source *j* to source *i*.
- *H*_1_: [**A**_*k*_]_*i,j*_ =0 for any *k* = 1, 2,⋯, *K*, i.e., there exists a causal influence from source *j* to source *i*.

Given that the VAR coefficients 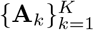 are unknown, to test this hypothesis, reliable estimates 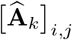, 1 ≤ *i, j* ≤ *M* and 1 ≤ *k* ≤ *K* are needed. However, such accurate estimates are often elusive due to limited observation horizon *T* compared to the number of parameters. Granger causality (Granger, 1969; Geweke, 1984, 1982) addresses this issue by considering the “bulk” effect of the VAR model coefficients through the prediction error metric. To this end, in assessing the causal influence of source *j* on source *i* two competing models are considered:

- *Full model*, where the activity of source *i* is modeled via the past activity of all the sources:

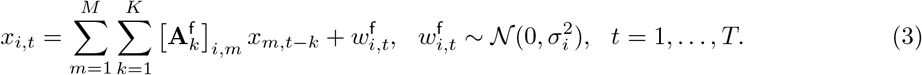
- *Reduced model*, where the contribution of the past of source *j* is removed from the full model by enforcing [**A**_*k*_]_*i,j*_ = 0, ∀*k* =1, 2,⋯, *K*:

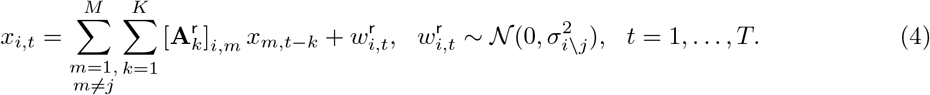

Note that we here use the *conditional* notion of Granger causality (Geweke, 1984), which includes all the processes *x_m_*,,, *m* ≢ *j* in both the reduced and full models. The process noise variables 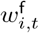 and 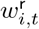 have different variances given by 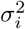 and 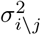, respectively. Define

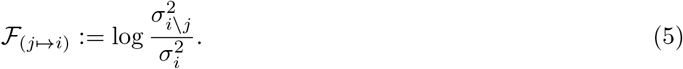

Clearly, when *j* has no causal influence on *i*, 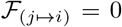, otherwise 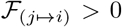, since the reduced model is nested in the full model, i.e., 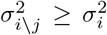. In practice, the VAR model coefficients 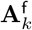 and 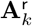, as well as the prediction variances 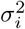 and 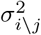 need to be estimated from the data. Let 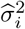 and 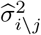 be the respective estimates of the prediction variances of the full and reduced models. Then, the resulting estimate 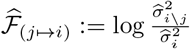 is a data-dependent random variable. Using 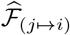, the previous hypotheses *H*_0_ and *H*_1_ for causality can be replaced by those of Granger causality (Greene, 2003):

- 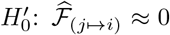, or equivalently 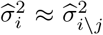. This implies that including the activity history of source *j* does not significantly improve the prediction error of source *i*, i.e., there is no Granger causal link from *j* to *i*.
- 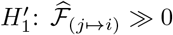, or equivalently 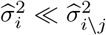. This implies that including the activity history of source *j* significantly improves the prediction accuracy of source *i*, i.e., there is a Granger causal link from *j* to *i*.

The test statistic 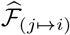 is referred to as the GC metric. In order to perform the latter hypothesis test, the asymptotic distribution of 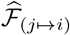 is utilized to obtain p-values (Kim et al., 2011). More specifically, under mild conditions, 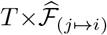 converges in distribution to a chi-square random variable with *K* degrees of freedom, i.e., *χ*^2^(*K*) (Wald, 1943; Davidson and Lever, 1970).

### 4.3. Challenges of GC Analysis for MEG

When it comes to GC analysis of cortical sources using MEG, there are several outstanding challenges:

1. *Indirect and Low-dimensional Sensor Measurements*. The foregoing notion of Granger causality assumes that the source time-series 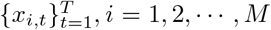 are directly observable. However, MEG only provides indirect and low-dimensional sensor measurements **y**_*t*_ ∈ ℝ^*N*^, where typically *N* ≪ *M*. As such, GC analysis of MEG data inherits the ill-posedness of estimating high-dimensional sources from low-dimensional sensor measurements (Wipf et al., 2010; Tait et al., 2021).
2. *Limited Observation Duration.* In order to obtain accurate estimates of the VAR model parameters and consequently prediction variances of the full and reduced models, typically observations with long duration *T* are required. However, the observation length is limited by the typically short duration of cognitive or sensory experimental trials. Even if trials with long duration were available, for the stationary model of Eq. (2) to be valid (i.e., static VAR parameters), *T* may not be chosen too long.
3. *Precise Statistical Characterization of the GC Links*. While the asymptotic distribution of the null hypothesis in the classical GC setting allows to obtain p-values, it is not clear how this asymptotic distribution behaves under the indirect and low-dimensional observations given by MEG. Furthermore, p-values only control Type I error, and in order to precisely characterize the statistical strength of the detected GC links, Type II errors need to also be quantified.

Existing methods aim at addressing the aforementioned challenges separately. In order to address challenge 1, source localization is used in a two-stage approach, where the cortical sources are first estimated using a source localization method, then followed by GC analysis (Cai et al., 2021, 2018; Owen et al., 2012); in order to address challenge 2, regularized least squares estimation is used to reduce the variance of the estimated VAR parameters (Endemann et al., 2022; Bolstad et al., 2011); and challenge 3 is usually addressed using non-parametric statistical testing, which may have limited power due to the large number of statistical comparisons involved (Cheung et al., 2010; Sekihara et al., 2010; Manomaisaowapak et al., 2021). It is noteworthy that these challenges are highly inter-dependent. For instance, the biases incurred by the source localization stage in favor of addressing challenge 1, may introduce undesired errors in the VAR parameter estimation to address challenge 2 (Schoffelen and Gross, 2009). Similarly, using regularized estimators to address challenge 2 introduces biased in the test statistics used in addressing challenge 3.

### 4.4. Proposed Solution: Network Localized Granger Causal (NLGC) Inference

We propose to address the foregoing challenges simultaneously and within a unified inference framework. To this end, we first cast Granger causal inference as an inverse problem using the generative models of Eqs. (2) and (1). To address the parameter estimation challenge of this inverse problem, we leverage sparse connectivity in cortical networks and utilize ℓ_1_-regularized estimation of the VAR parameters. Finally, to characterize the statistical strengths of the identified GC links, we establish the asymptotic properties of a test statistic, namely the de-biased deviance difference, which will allow us to parametrically quantify both Type I and Type II errors rates and also control the false discovery rate. We refer to our proposed method as the Network Localized Granger Causality (NLGC) analysis. The main building blocks of NLGC are introduced in the remaining part of this subsection.

#### 4.4.1. Efficient Parameter Estimation and Likelihood Computation

It is straightforward to show that this classical GC metric, i.e., log-ratio of the prediction variances of the reduced and full models in Eq. (5) is equivalent to the difference of the log-likelihoods of the full and reduced models, for linear Gaussian generative models. This correspondence has led to the generalization of the GC metric to non-linear and non-Gaussian settings (Kim et al., 2011; Sheikhattar et al., 2018).

We take a similar approach to generalize the classical notion of GC for direct observations of the sources to our indirect observations given by the MEG sensors. Recall that for assessing the GC from source *j* to *i*, we considered the full and reduced models given by Eqs. (3) and (4). Let 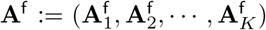 and 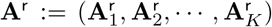 be the VAR parameters matrices, and 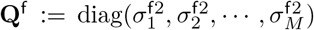 and 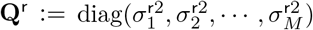 be the process noise covariance matrices of the full and reduced models, respectively. The main difference between these sets of parameters is that 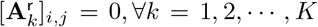. Let the log-likelihoods of the MEG observations under the full and reduced models be defined as:

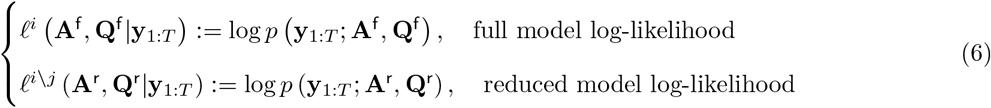

Let 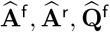, and 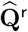 be the regularized maximum likelihood estimates of the corresponding parameters. We then define the GC metric from source *j* to *i* given the MEG observations as (Kim et al., 2011; Sheikhattar et al., 2018; Soleimani et al., 2020):

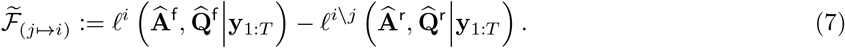

As for the regularization scheme, we consider ℓ_1_-norm regularized maximum likelihood estimation. Let **a**_*i*_ be the *i*^th^ row of **A**, correspond to all the network interactions towards source *i*. The parameters are estimated as:

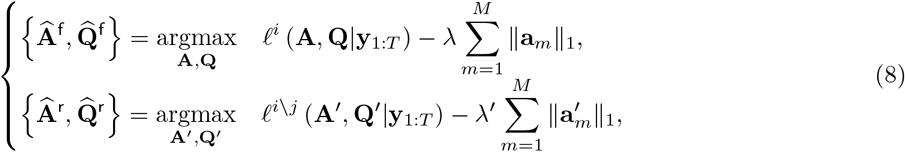

where λ, λ′ are regularization parameters that are tuned in a data-driven fashion using cross-validation (See Appendix A.1 for details). Since the source activity 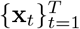 is not directly observable, we employ an instance of Expectation-Maximization (EM) algorithm (Shumway and Stoffer, 1982; Dempster et al., 1977) to solve the regularized maximum likelihood problem. The EM algorithm is an iterative procedure which maximizes a lower bound on the log-likelihood function and provides a sequence of improving solutions. The EM algorithm has two steps: 1) The Expectation step (E-step) where we calculate the expectation of the log-likelihood of both the observed and unobserved variables given the observations and a current estimate of the parameters to construct a lower bound on the actual observation log-likelihood, and 2) The Maximization step (M-step) where we maximize the surrogate function obtained in the E-step to update the estimate of the unknown parameters.

More specifically, we illustrate these two steps for estimating the parameters of the full model; the case of reduced model is treated in a similar fashion. Let the unknown parameters be denoted by ***θ*** := (***θ***_1_,…, ***θ***_*M*_), where 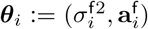 is the corresponding unknown parameters of the *i*^th^ source with 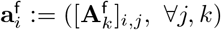. The EM algorithm in this case comprises the following steps:

##### The E-step

Starting from an initial point, let us denote the parameter estimates at the *l*^th^ iteration of the EM algorithm by 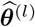. At the E-step, we define the so-called Q-function:

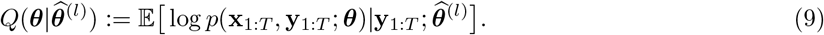

Given the linear Gaussian state-space model used as our generative model, the expectation in Eq. (9) requires the first and second moments of **x**_*t*_ given **y**_1:*T*_ and 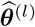 and can thus be efficiently computed using Fixed Interval Smoothing (FIS) (Anderson and Moore, 2005).

##### The M-step

At the M-step, we update the parameters as

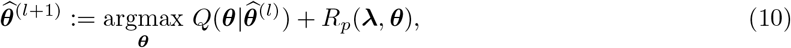

where *R_p_*(**λ**, ***θ***) is a regularization function to enforce sparsity of the parameters. Here, we use the FASTA algorithm to solve the optimization problem in Eq. (10) (Goldstein et al., 2014). These steps continue until convergence of the iterates 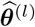. To assess convergence, the log-likelihood of the MEG observations is calculated (Gupta and Mehra, 1974) at each iteration, to check whether the successive improvements of the log-likelihood fall below a specified threshold. Fig. 8 gives an overview of the EM algorithm, which is derived in full details in Appendix A.

**Figure 8:**
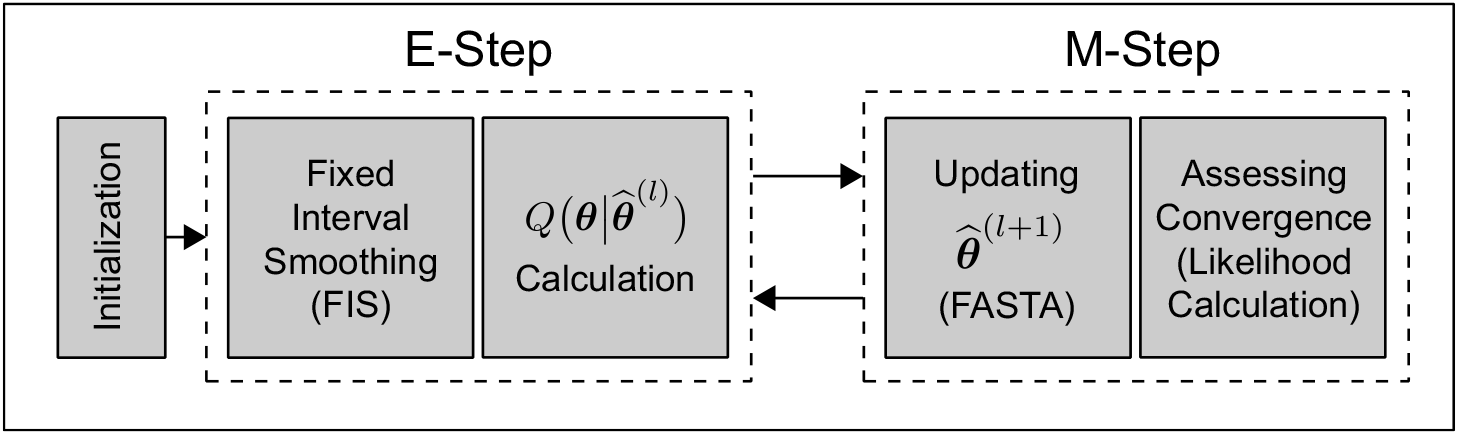
Block diagram of the EM algorithm for sparse VAR parameter estimation.

Employing the foregoing EM procedure, one can reliably estimate the set of parameters ***θ*** corresponding to the full model and reduced models for all possible links (*j* ↦ *i*) and evaluate the log-likelihoods to form the GC metric 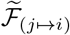 of Eq. (7), for all *i, j* = 1, 2,⋯, *M*, *i* ≠ *j*.

#### 4.4.2. Computational Complexity of the Parameter Estimation Procedure

Applied to MEG, off-the-shelf solvers do not scale well with the dimensions of the source space *M*, sensor space *N*, and observation length *T*. We employ several solutions to address this need for scalability of the parameter estimation procedure:

1. First, we use a low-rank approximation to the lead-field matrix that reduces the effective dimensionality of the source space. This approach is explained in detail in Section 4.5.1.
2. We use the steady-state solution to the smoothing covariance matrices involved in FIS that notably speed up the computations. This approach is explained in detail in Appendix A.2.
3. We use the Fast Adaptive Shrinkage/Thresholding Algorithm (FASTA) algorithm to efficiently solve the *ℓ*_1_-regularized optimization in the M-step. This approach is explained in Appendix A.1.
4. We efficiently evaluate the various log-likelihood functions, which are key for cross-validation and the EM stopping criterion, using the innovation form of the smoothed states (Gupta and Mehra, 1974).

In what follows, we discuss the implications of these algorithmic solutions in reducing the computational complexity of our EM-based parameter estimation procedure used for solving Eq. (8), in comparison to existing work.

As it will be shown in Section 4.5.1, Solution (1) results in an effective lead-field matrix with *rM* columns, where *M* is the number of cortical patches used and *r* ≥ 1 is the number eigenmodes retained in the low-rank representation of the lead-fields in each patch. Also, Solution (2), using the steady-stake Kalman filtering/smoothing, reduces the total number of state covariance matrix inversions in the FIS procedure from *T* to 2, by only adding 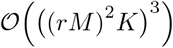 multiplications required to find the steady-state covariance matrices (Malik et al., 2010). Considering the cubic dependence of matrix inversion to the matrix dimension, each instance of FIS requires 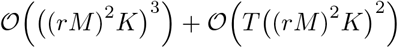 multiplications, which can then be used to form the elements of the Q-function in the E-step.

At the M-step, Solution (3) uses FASTA to update the parameters. As a gradient-based method, for an optimality gap of *ε* > 0, it requires 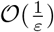 iterations, and each iteration requires 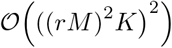 multiplications (Beck and Teboulle, 2009; Goldstein et al., 2014). Here, we denote the complexity of FASTA by 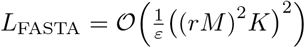. Next, Solution (4) provides an efficient method to compute the log-likelihood of the MEG observations (Gupta and Mehra, 1974), which only includes matrix additions and matrix by vector multiplications based on the quantities already calculated at the FIS procedure, adding up to 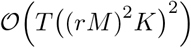 multiplications. Finally, letting *L*_EM_ be the number of EM iterations, each application of the EM algorithm requires 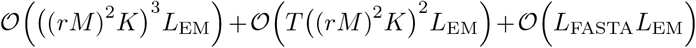 multiplications.

The problems in Eq. (8) need to be solved for both the full and reduced models. The only difference between the full model and reduced model corresponding to the link (*j* ↦ *i*) is the fact that in the reduced model, one set of the cross-coupling coefficients **a**_*i,j,k*_ (*k* = 1,⋯, *K*) are constrained to be zero during the EM procedure (See Remark 2 in Appendix A.1). The total number of such estimation problems to be solved is 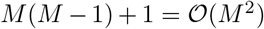. Thus, the overall computational complexity of our parameter estimation procedure is given by 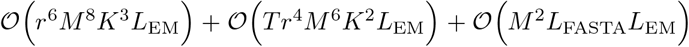. In the applications of interest in this work, typically the convergence criteria is satisfied with a choice of *L*_FASTA_ ≈ 100 and *L*_EM_ ≈ 1000, which mitigates the dependence of the overall computational complexity on these parameters.

The improvements achieved by Solutions (1) and (2) provide notable computational savings over existing work (Nalatore et al., 2009; Cheung et al., 2010; Sekihara et al., 2010; Long et al., 2011; Lamus et al., 2012): 1) If the low-rank approximation to the lead-field matrix is not used, the term *r* is replaced by 61 (see Section 4.5.1 for details). Given that we use a value of *r* = 4 in our work, this amounts to a ~ 10^7^-fold reduction in the complexity of the leading term that is 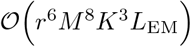; 2) If the steady-state filtering/smoothing is not used, the first term in the computational complexity of the EM procedure would be increased to 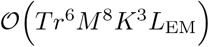. Our approach reduces this term by a factor of *T*, which in the applications of interest in this paper amounts to a ~ 10^3^-fold reduction in complexity.

#### 4.4.3. Statistical Test Strength Characterization

The next component of NLGC is the characterization of the statistical significance of the obtained GC metrics. Let 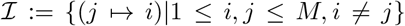 be the set of all possible GC links among *M* sources. Consider the link 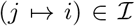 and let us represent the corresponding parameters of the full and reduced models of the link as ***θ***^f^ and ***θ***^r^, respectively, where for ***θ***^r^ we have 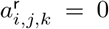, ∀*k*. It is worth noting that the number of parameters to be estimated in the full and reduced models are *M*^f^ := *K*(*rM*)^2^ and *M*^r^ := *K*(*rM*)^2^ – *Kr*^2^, respectively. We define the null hypothesis *H*_(*j*↦*i*),0_ : ***θ*** = ***θ***^r^ for the case that no GC link exists, and the alternative *H*_(*j*↦*i*),1_ : ***θ*** = ***θ***^f^ for the existence of a GC link from source *j* to source *i*. A conventional statistic for testing the alternative against the null hypothesis is the *deviance difference* between the estimated full and reduced models defined as

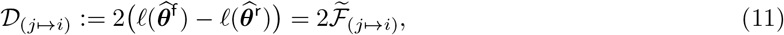

where ℓ(***θ***) := log*p*(**y**_1:*T*_; ***θ***) is the log-likelihood of the observations. Large values of 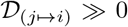 indicate a large improvement in the log-likelihood of the full model compared to that of the reduced model, which implies the existence of a GC link. Similarly, 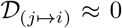 can be interpreted as the absence of a GC link from source *j* to source *i* (Kim et al., 2011).

Conventionally, the asymptotic distribution of the deviance difference is derived as a chi-square distribution, thanks to the asymptotic normality of maximum likelihood estimators (Wald, 1943; Davidson and Lever, 1970). However, due to the biases incurred by ℓ_1_-norm regularization, the estimates are no longer asymptotically normal. To remove the bias and obtain a statistic with well-defined asymptotic behavior, we use the de-biased version of the deviance difference introduced in (Sheikhattar et al., 2018; Soleimani et al., 2020):

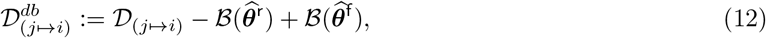

where 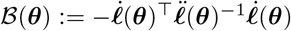 is the empirical bias incurred by ℓ_1_-norm regularization (van de Geer et al., 2014), with 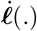 and 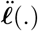 denoting the gradient vector and Hessian matrix of the log-likelihood function ℓ(.), respectively. Removal of the bias allows to recover the well-known asymptotic behavior of the deviance difference. We characterize these distributions using the following theorem:

#### Theorem 1.

*The de-biased deviance difference defined in* Eq. (12) *converge weakly to the following distributions, under the null and alternative hypotheses (as T* ↦ ∞):

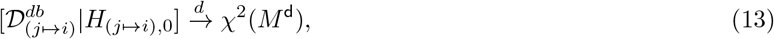

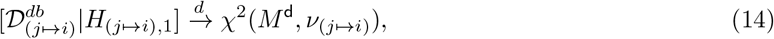

*where χ*^2^(*q*) *denotes the central chi-square distribution with q degrees of freedom, and χ*^2^ (*q, ν*) *represents the non-central chi-square distribution with q degrees of freedom and non-centrality parameter ν, with M*^d^ := *M*^f^ – *M*^r^ = *K r*^2^.

*Proof.* See Appendix B.

In words, Theorem 1 states that the asymptotic distribution of the de-biased deviance difference in the absence and presence of a GC link is distributed according to central and non-central χ^2^ distributions, both with degree of freedom *Kr*^2^, i.e., the number of VAR parameters from patch *j* to *i*, respectively. The non-centrality parameter in Eq. (14) can be estimated as 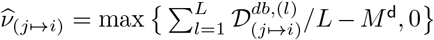 where 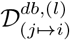 is the *l*^th^ sample of the de-biased deviance computed from *L* ≥ 1 independent trials (Saxena and Alam, 1982). We will next show how the result of Theorem 1 can be used for FDR control as well as characterizing the test strength.

##### FDR control

Recall that rejection of the null hypothesis for a given source and target pair implies the existence of a GC link. As a consequence, determining GC links among the source and target pairs requires preforming *M*(*M* – 1) multiple comparisons, which may result in high false discovery. To address this issue, we employ the Benjamini-Yekutieli (BY) FDR control procedure (Benjamini and Yekutieli, 2001). Consider the link 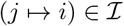. According to the first part of Theorem 1, if the null hypothesis is true, i.e., the GC link does not exist, the corresponding de-biased deviance difference is central chi-square distributed. Thus, at a confidence level 1 – *α*, the null hypothesis *H*_(*j*↦*j*),0_ is rejected if 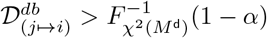 where 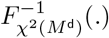 is the inverse cumulative distribution function (CDF) of the central χ^2^ distribution with *M*^d^ degrees of freedom. Using the BY procedure, the average FDR can be controlled at a rate of 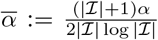 where 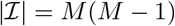 represents the cardinality of the set 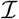.

**Algorithm 1.**
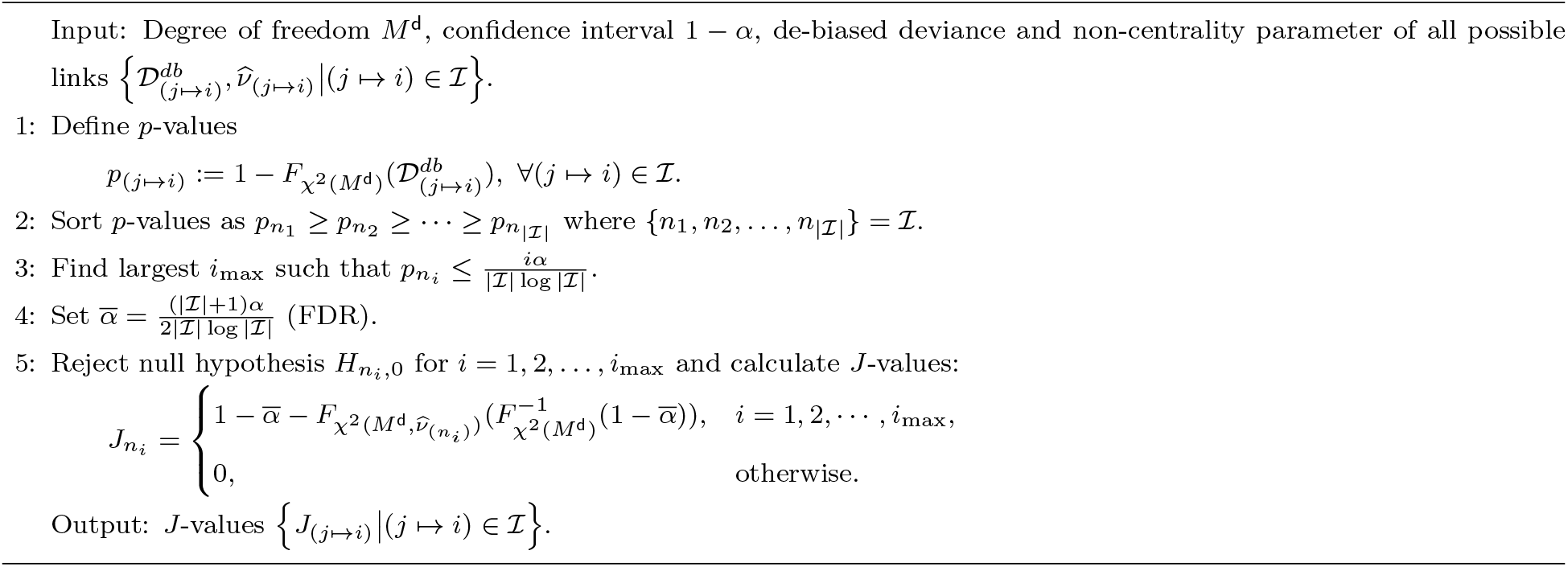
FDR control and test strength characterization

##### Test Strength Characterization

To determine the test strength, we use the second part of Theorem 1 as well to quantify Type II errors. To this end, the false negative rate at the given confidence level 1 – *α* for a source-target pair (*j* ↦ *i*) is given by 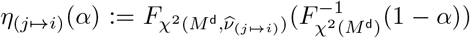 where 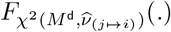 denotes the non-central χ^2^ distribution with *M*^d^ degrees of freedom and non-centrality parameter 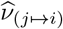. Given the false negative rate, we use the Youden’s *J*-statistic (Youden, 1950) to summarize the strength of the test as:

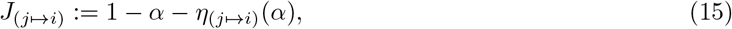

for the given confidence level 1 – *α*. The *J*-statistic has a value in the interval [0,1] summarizing the performance of a diagnostic test. When *J*_(*j*↦*i*)_ ≈ 0, the evidence to choose the alternative over the null hypothesis is weak, i.e., the GC link is likely to be missing. On the other hand, when *J*_(*j*↦*i*)_ ≈ 1, both the false positive and negative rates are close to zero, implying high test strength, i.e., strong evidence in support of the GC link.

The overall statistical inference framework is summarized in Algorithm 1. Finally, obtaining the *J*-statistics for all links, we can construct the GC map **Φ** as follows

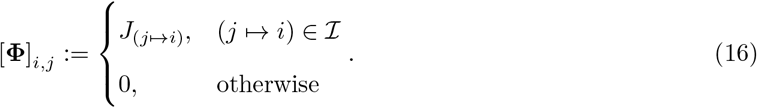

It is worth noting that to repeatedly evaluate the de-biased deviance difference statistic, one needs to efficiently calculate the log-likelihood function ℓ(.), which is done using the innovation form described in (Gupta and Mehra, 1974). In the spirit of easing reproducibility, a python implementation of the NLGC is available on the open source repository Github (Soleimani and Das, 2022).

### 4.5. Dimensionality Reduction and VAR Model Order Selection

There are two remaining ingredients of NLGC which are key to ensure its scalability, namely, reducing the dimensionality of the source space and VAR model order selection.

#### 4.5.1. Source Space Construction and Eigenmode Decomposition

In practice, using MR scans of the participants, individual head models can be numerically computed and co-registered to each individual’s head using the digitized head shapes. We first define a cortical surface mesh-based source space for the ‘fsaverage’ head model (Dale et al., 1999), named ico-4, with average spacing of ~ 6 mm between any two neighboring sources, which is then morphed to each participant’s head model. The lead-field matrix is obtained by placing 3 virtual dipoles at each of the 5124 vertices of ico-4 source space and solving Maxwell’s equations. We further restrict the dipoles to be normal to the cortical surface, so that the resulting lead-field matrix **C** has *M* = 5124 columns of length *N* each (Gramfort et al., 2013a, 2014). Solving the NLGC inverse problem over this source space is quite computationally demanding, as the computational time of FIS scales as 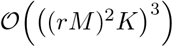 (See Section 4.4.2). We thus need to reduce the dimension of the lead-field matrix to control the computational complexity.

To this end, we summarize the contribution of the dipoles placed on the ico-4 source space vertices within a given region using their principal components (Limpiti et al., 2006; Cheung et al., 2010). We start from a coarse surface mesh-based source space, namely ico-1, with 84 vertices (42 vertices per hemisphere). We consider the Voronoi regions based on the geodesic distance between these vertices induced by ico-1 vertices over the original ico-4 vertices, so that all the ico-4 vertices are partitioned into 84 non-overlapping patches (Babadi et al., 2014). The Voronoi regions around each of the ico-1 vertices are referred to as *cortical patches* in this work. We then approximate the contribution of the dipoles placed on the ico-4 vertices within each cortical patch by the first *r* leading eigenvectors of the partial lead-field matrix following singular value decomposition (SVD). We refer to these leading eigenvectors as *eigenmodes*. As such, the number of columns in the *effective* lead-field matrix is reduced to *r* × 84, as opposed to original 5124, which significantly reduces the computational complexity. In addition to providing computational savings, dimensionality reduction through retaining the leading eigenmodes of the lead-field sub-matrices serves as denoising by suppressing the effect of small lead-field errors (which are expected to appear in eigenmodes with small singular values).

Fig. 9 shows a schematic depiction of the eigenmode decomposition for a given patch with *r* = 2 eigenmodes. For this example, the 10 × 7 lead-field matrix of the cortical patch is reduced to a 10 × 2 matrix, for which the two eigenmodes capture the main contributions of the patch to the MEG sensors. In other words, we summarize all the dipoles placed on ico-4 vertices within each cortical patch by the best *r effective* dipoles, which explain most of the lead-field variance within that cortical patch. With increasing r, the approximation gets better in a similar way that a finer cortical mesh improves cortical current density approximation. The parameter *r* can be chosen by controlling the reconstruction error at a desired level. We will provide an example of this choice in the following subsection.

**Figure 9:**
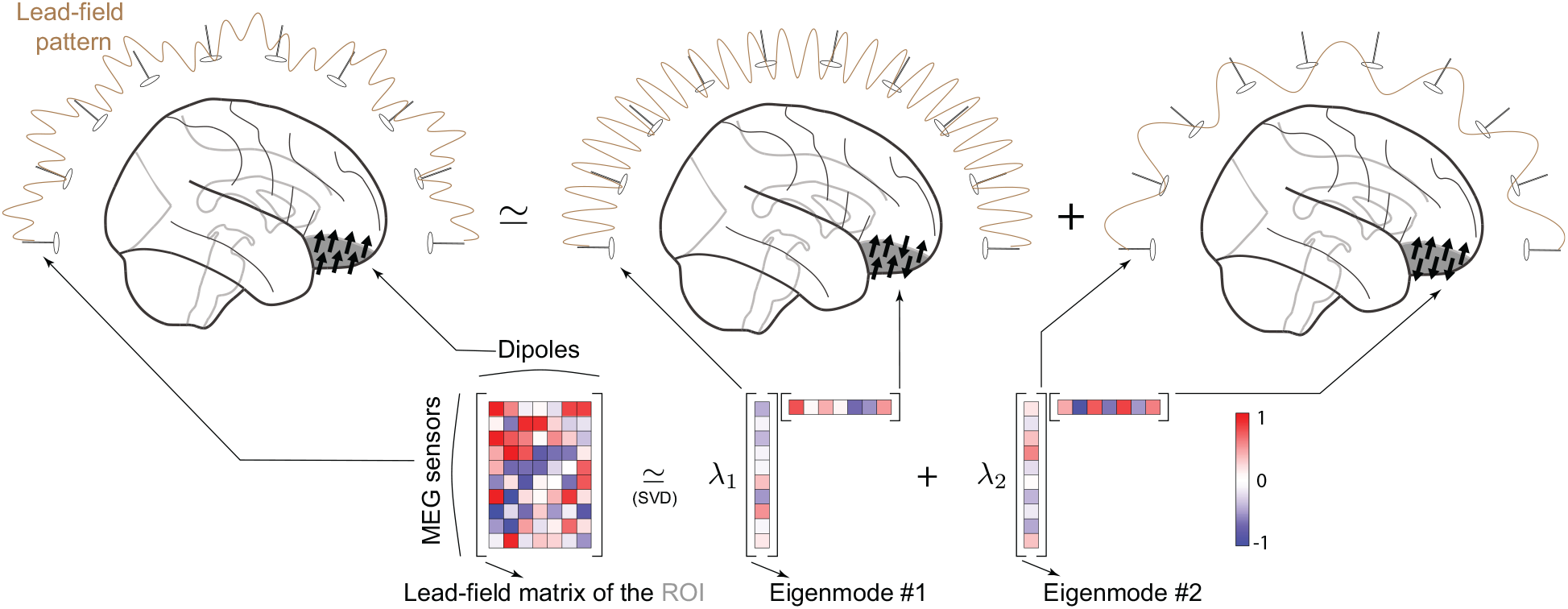
An illustration of low-rank approximation to the lead-field matrix using eigenmode decomposition using *r* = 2 eigenmodes. The contribution of the 7 dipoles to 10 MEG sensors is originally captured by a 10 × 7 submatrix of the lead-field matrix (left), whereas using the eigenmode decomposition, it can be approximated by two 10-dimensional eigenmodes (right), resulting in a 10 × 2 effect sub-matrix.

#### 4.5.2. VAR Model Order Selection

In Section 4.4, the VAR model order *K* is assumed to be known. To estimate *K* in a data-driven fashion, we utilize the *Akaike Information Criterion* (AIC) to determine which model order best fits the MEG observations (Ding et al., 2018). Given a set of candidate model orders 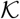 for *K*, the optimal model order can be chosen as:

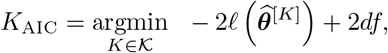

where *df* is the degrees of freedom of the ℓ_1_-norm regularized maximum likelihood problem (Zou et al., 2007) and 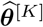 denotes the estimated parameters corresponding to a VAR(*K*) model.

Ideally, one can search within a large set of candidate values for *K* and *r* (number of eigenmodes) and choose the optimal pair according to an information criterion (Ding et al., 2018). However, due to high computational complexity of the estimation procedure in NLGC, especially for higher values of *K* and *r*, we first pick a suitable value for the number of eigenmodes *r*, followed by choosing the VAR model order *K* via AIC.

To choose *r*, we require that at least 85% of the variance within each ROI can be explained using *r* eigenmodes. Depending on the subject’s head model and also the location of the dipoles, the choice of *r* may vary. For the MEG data in this study, *r* = 4 eigenmodes sufficed to capture at least 85% of the variance. Fig. 10A shows the histogram of explained variance ratio for all ROIs using *r* = 2, 3, 4 eigenmodes corresponding to 3 representative subjects.

**Figure 10:**
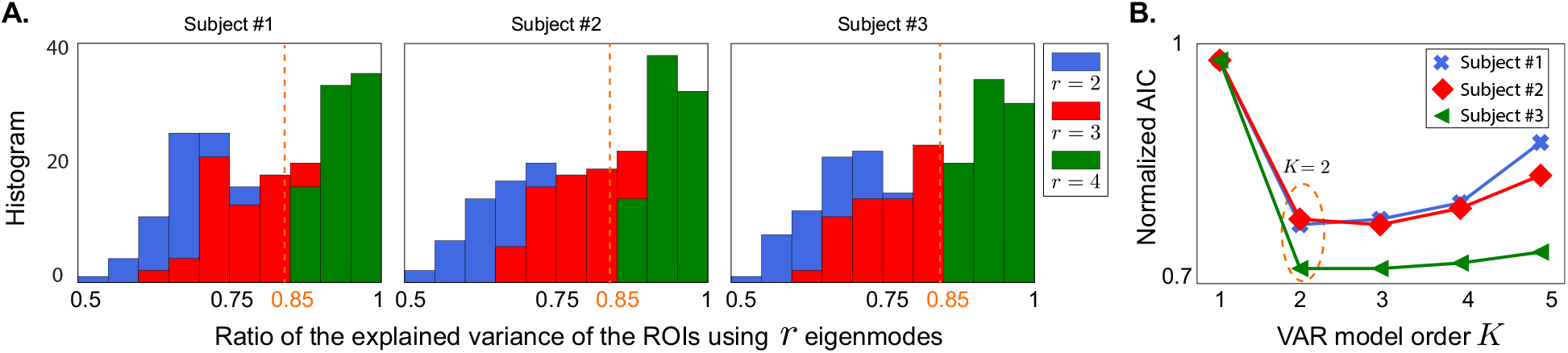
Model selection curves. **A.** Histogram of the ratio of the explained variance to total variance for all ROIs using *r* = 2, 3,4 eigenmodes for head models of three representative subjects. With *r* = 4 eigenmodes, at least 85% of the variance can be explained for all ROIs. B. AIC curve for *r* = 4 egienmodes, suggesting a choice of *K* = 2 for the VAR model order for the three representative subjects.

Once *r* = 4 is fixed, we use AIC to pick the optimal value of *K*. For the MEG data in this study, *K* = 2 was the optimal choice according to AIC for all subjects. Fig. 10B shows the AIC curves of the same 3 subjects as in panel A. Even though in some cases (e.g. subject 2), a choice of *K* = 3 results in a slight improvement compared to *K* = 2, to reduce the overall run-time of our inference framework, we picked *K* = 2 for all cases.

### 4.6. MEG Experiments: Procedures and Recordings

The data analyzed in this study was a part of a larger experiment whose results will be reported separately. Out of 36 total participants who completed the MEG experiment, 24 participants completed the structural MRI scans. Additionally, 2 subjects were excluded due to bad fiducials measurements. Ultimately, 22 subjects, 13 younger adults (5 males; mean age 21.1 years, range 17–26 years) and 9 older adults (3 males; mean age 69.6 years, range 66-78 years) were included in the analysis. All participants had clinically normal hearing (125–4000 Hz, hearing level ≤ 25 dB) and no history of neurological disorder.

The study was approved by the University of Maryland’s Institutional Review Board. All participants gave written informed consent and were compensated for their time. Subjects came in on two different days. MEG auditory task recording was performed on the first day and structural MRIs were scanned on the second day. Neural magnetic signals were recorded in a dimly lit, magnetically shielded room with 160 axial gradiometer whole head MEG system (KIT, Kanazawa, Japan) at the Maryland Neuroimaging Center. The MEG data were sampled at 2 kHz, low pass filtered at 200 Hz and notch filtered at 60 Hz. Participants laid supine position during the MEG experiment while their head was in the helmet and as close as possible to the sensors. The head position was tracked at the start and end of the experiment with 5 fiducial coils. During the task subjects were asked to stare at the center of the screen and minimize the body movements as much as possible.

The resting state data were recorded before and after the main auditory task, each 90 s long in duration. During the resting state subjects fixated at a red cross at the center of grey screen. 100 repetitions of 500 Hz tone pips were presented at the end. During the tone pips task, subjects were staring at a face image at the center of screen and were asked to silently count the number of tone pips. The tones were presented at a duration of 400 ms with a variable interstimulus interval (1400 ms, 1200 ms, 1000 ms). The tone pip task was around 150 s long and was divided into two trials, 40 s after the beginning of the first tone pip onset resulting in two trials. In summary, we analyzed the GC link counts in resting state and listening to tone pips task, each consisted of two trials.

### 4.7. Pre-processing and Data Cleaning

All the pre-processing procedures have been carried out using `MNE-python 0.21.0` (Gramfort et al., 2013a, 2014). After removing the noisy channels, temporal signal space separation (tsss) was used to remove the artifacts (Taulu and Simola, 2006). The data were filtered between 0.1 Hz and 100 Hz using a causal FIR filter (with `phase=‘minimum’` setting). Independent component analysis (extended Infomax algorithm, with `method=‘infomax’` and `fit_params=dict(extended=True)` settings) was applied to extract and remove cardiac and muscle artifacts (Bell and Sejnowski, 1995; Lee et al., 1999). The initial 5 seconds of the data were removed and the subsequent 40 seconds were extracted. Finally, the data were filtered to the desired frequency bands using causal FIR filters followed by downsampling to 50 Hz.

### 4.8. NLGC Parameter Settings

As mentioned in Section 4.5.2, the VAR model order *K* is selected via AIC over a set of candidates 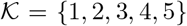. The regularization parameter for the ℓ_1_-norm are chosen using a standard 5-fold cross-validation over the range [10^-15^,1] spanned by 25 logarithmically-spaced points (Appendix A.1, Remark 3). As for the convergence of the EM algorithm, we used a normalized error tolerance of tol = 10^-5^, with a maximum number of 1000 iterations (Algorithm 2). For all simulation studies as well as real data analysis FDR was controlled at 0.1% using the BY procedure.

#### 4.8.1. Parameters for the Illustrative Example

We considered *M* = 84 cortical patches, whose activities are projected onto the MEG sensor space with N = 155 sensors. We simulated 3 different realizations (with *T* = 1000 samples each) for each run. To simplify the projection onto the MEG sensors, we considered a single lead-field vector for each cortical patch, generated via drawing 155 independent samples from a standard normal distribution. This simplification using a single lead-field vector per patch could be thought of as taking a random linear combination of all the lead-field vectors within a cortical patch as the representative of its activity. The noise measurement covariance matrix was assumed to be diagonal **R** = *σ*^2^**I** where *σ*^2^ was chosen to set the SNR at 0 dB. The cortical patch activities were simulated as a VAR(5) process. Among them, 8 patches were randomly selected to carry the dominant activities, i.e., explaining 90% of the total signal power. To compare the performance of NLGC with a two-stage method using MNE, we first obtained the source estimates for the first stage as:

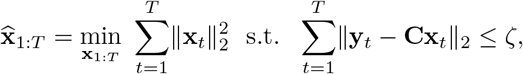

for some *ζ* > 0. Given the source estimates, we then fit the VAR models to obtain the network parameters (Appendix A.3). Then, the same statistical inference framework used in NLGC was applied to extract the GC links in the second stage.

#### 4.8.2. Parameters for the Simulated MEG Data Using a Head-Based Model

We computed the forward solution for ico-4 source space from a representative younger subject’s head model via MNE-python 0.21.0 and then obtained the low-rank lead-field matrix approximation over ico-1 source space using the previously mentioned dimensionality reduction strategy (see Section 4.5.1 for details). Each of the cortical patches corresponding to ico-1 vertices had *r*_gen_. eigenmodes, resulting in 84 × *r*_gen_. lead-field columns, which are summarizing the contribution of 5124 ico-4 sources, partitioned into 84 groups according to the Voronoi regions formed over the cortical manifold. As a result, in the generative model, the lead-field matrix has *M* = 84 × *r*_gen_. columns and *N* = 155 rows. The dipole activities 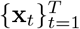 were generated using VAR(3) processes with *T* = 3000 time points (3 segments, 1000 samples each). With 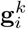 denoting the *k*^th^ eigenmode of the *i*^th^ cortical patch, the MEG observation at time *t* is generated as

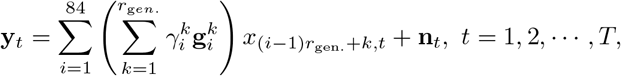

where 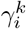 are drawn uniformly in the interval [−1, 1] and **n**_*t*_ is a zero mean Gaussian random vector with a diagonal covariance matrix **R** = *σ*^2^**I**. The value of *σ*^2^ is determined according to the desired SNR level which is set to 0 dB, unless otherwise stated.

We considered varying numbers of dominant cortical patches, *m* = 2, 4,⋯, 20 that explain 90% of the total signal power. The remaining 10% of the signal power was uniformly distributed as white noise among the rest of cortical patches. The true underlying GC network structure among the dominant cortical patches was assumed to have 20% sparsity, i.e., with *m* active cortical patches, there are ⌈0.2*m*(*m*–1)⌉ true GC links, where ⌈*z*⌉ denotes the smallest integer greater than or equal to *z*. For each *m*, we generated 10 different trials of the VAR processes, while randomly selecting cortical patches from the temporal and frontal lobes for each trial.

In all the four cases considered to assess the robustness of the algorithms, we used *r*_est_. = 2. To induce source model mismatch, we simply used *r*_gen_. = 10 (> *r*_est_.) eigenmodes for the data generation process. We also considered a relaxed link localization criterion in addition to the exact link localization criterion. The rationale behind the relaxed link localization criterion is as follows: Let (*j* ↦ *i*) be a true GC link, and let *N*(*i*) denote the 6 nearest cortical patches to cortical patch *i* over the ico-1 source space. If instead the link (*j*′ ↦ *i*′) is detected, we consider it a hit if *i*′ ∈ *N*(*i*) and *j*′ ∈ *N*(*j*). This way, we account for minor spatial localization errors. Note that in the exact link localization criterion, the link (*j* ↦ *i*) is considered a hit only if it is exactly detected by NLGC.

The NLGC settings were the same in all the aforementioned cases. For the two-stage methods, we used the standard MNE and dSPM methods as well as the Champagne algorithm implemented in MNE-python 0.21.0 using their default settings to localize the simulated MEG data into cortical time-courses. For each value of m, we ran NLGC and the three two-stage procedures and evaluated the performance of each method by calculating the hit rate (number of true detected links normalized by the total number of true links) and false alarm rate (number of spurious links normalized by the total number of non-GC links), both averaged over the 10 trials.

#### 4.8.3. Parameters in the Analysis of Experimentally Recorded MEG Data

For the MEG data that were recorded during an auditory task, we analyzed the connectivity between ROIs in frontal, temporal, and parietal lobes (in both hemispheres) that broadly comprise the auditory cortex, the fronto-parietal network, the cingulo-opercular network, the ventral attention network, and the default mode network, which are known to fluctuate with task versus rest conditions (Fox et al., 2005) and with aging (Kuchinsky and Vaden, 2020). The included ROIs are selected from the 68 anatomical ROIs in the Desikan-Killiany atlas (Desikan et al., 2006):

- Frontal: ‘rostralmiddlefrontal’, ‘caudalmiddlefrontal’, ‘parsopercularis’, ‘parstriangularis’.
- Temporal: ‘superiortemporal’, ‘middletemporal’, ‘transversetemporal’.
- Parietal: ‘inferiorparietal’, ‘posteriorcingulate’.

We then mapped the 84 cortical patches onto these 68 anatomical ROIs. To illustrate this procedure, consider the example given in Fig. 11. There are three representative cortical patches, denoted by *d_k_, k* = 1, 2, 3 with corresponding vertices in ico-1 (crosses) and ico-4 (arrows) mesh are shown with the same color.

**Figure 11:**
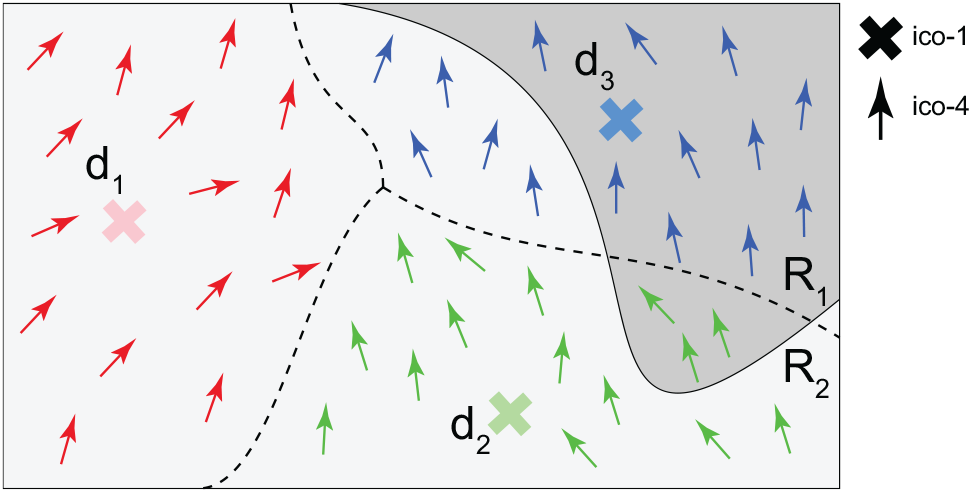
Illustration of anatomical ROI to cortical patch assignment. Three ico-1 vertices shown as *d*_1_ (red ×), *d*_2_ (green ×) and *d*_3_ (blue ×) as well as the corresponding ico-4 vertices (colored arrows) in the respective patches are shown with the same color coding. Two anatomical ROIs *R*_1_ (dark grey) and *R*_2_ (light gray) are also highlighted. Using the proposed association scheme, each cortical patch is assigned a pair of weights indicating its relative overlap with the two ROIs. Here, the association weights of *d*_1_, *d*_2_ and *d*_3_ are given by (0,1), (0.2, 0.8) and (0.67, 0.33), respectively.

The goal is to allocate the representative cortical patches between the two ROIs marked by *R*_1_ and *R*_2_. For each representative cortical patch, we compare the ratio of the number of ico-4 vertices that lie within each ROI and use it as an association weight between the representative cortical patch and the ROI. For the given example in Fig. 11, the association weights to *R*_1_ and *R*_2_ for the three representative cortical patches *d*_1_, *d*_2_, *d*_3_ are given by (0,1), (0.2, 0.8), and (0.67, 0.33), respectively. Using this many-to-one mapping, the obtained NLGC map **Φ**, which represents the GC links among the ico-1 cortical patches, can be translated into a connectivity map among the 68 ROIs as follows. Let **W** ∈ ℝ^84×68^ denote the aforementioned association weight matrix, where [**W**]_*i,j*_ is the association weight of the *i*^th^ representative cortical patch to the *j*^th^ ROI. The transformed connectivity map 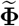 is then defined as 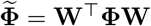.

As an example of this transformation, consider the setting of Fig. 11 and suppose that NLGC only detects one GC link (*d*_2_ ↦ *d*_2_). Assuming that there are only 3 patches *d*_1_, *d*_2_, and *d*_3_ in the model, we have:

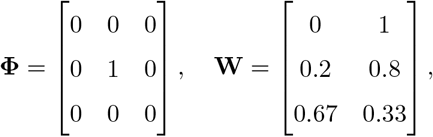

where the weight matrix **W** contains the association weights of the setting in Fig. 11. The transformed connectivity matrix is thus given by:

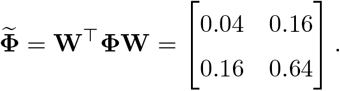

We can then interpret 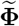 as follows: the captured link (*d*_2_ ↦ *d*_2_) is decomposed into several possible links between the 2 anatomical ROIs *R*_1_ and *R*_2_, namely (*R*_1_ ↦ *R*_1_) with a weight of 0.04, (*R*_1_ ↦ *R*_2_) with a weight of 0.16, (*R*_2_ ↦ *R*_1_) with a weight of 0.16, and (*R*_2_ ↦ *R*_2_) with a weight of 0.64. Notably, the elements of 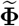 add up to one, which guarantees that the link (*d*_2_ ↦ *d*_2_) is not double-counted under the many-to-one mapping from the patches to anatomical ROIs, and thus the total number of GC links is preserved.

The VAR model order and the number of eigenmodes are chosen as *K* = 2 and *r* = 4 using AIC criterion. The details of the model selection is described in Section 4.5.2. To obtain the directed networks between frontal, temporal, and parietal areas, for each of the Delta+Theta and Beta frequency bands of interest, we encoded the inferred connectivity maps for each subject in each trial and condition using a 9-dimensional vector, where each entry represented the number of detected GC links corresponding to the connectivity types *A* ↦ *B* where *A, B* ∈ {Frontal, Temporal, Parietal}. For the inter- vs. intra-hemispheric refinement of our analysis, encoded the GC maps using a 36-dimensional vector in which the entries also distinguished between the connectivity across and within hemispheres, i.e., *A*(*h*) ↦ *B*(*h*) where *h* ∈ {left hemisphere, right hemisphere} and *A, B* ∈ {Frontal, Temporal, Parietal}.

Another key parameter that may affect the performance of NLGC is the choice of the trial duration *T*. To investigate the effect of the trial duration on the performance of NLGC, we repeated NLGC analysis using different values of *T* corresponding to the first 20, 25,⋯, 40 seconds of the data. The results corresponding to the younger participants under the tone processing condition over the Delta+Theta band is shown are Fig. 12. As it can be observed from the figure, for small values of *T* the detected networks are quite sparse, as the algorithm does not have enough statistical power to detect all relevant links. It is worth noting that NLGC did not capture any GC links using only the first 10 seconds of the data. For ~ 30 s and higher, the captured GC network stabilizes and converges. Therefore, the choice of 40 s used in our analysis is taken conservatively to make sure that enough data points are available for GC link detection.

**Figure 12:**
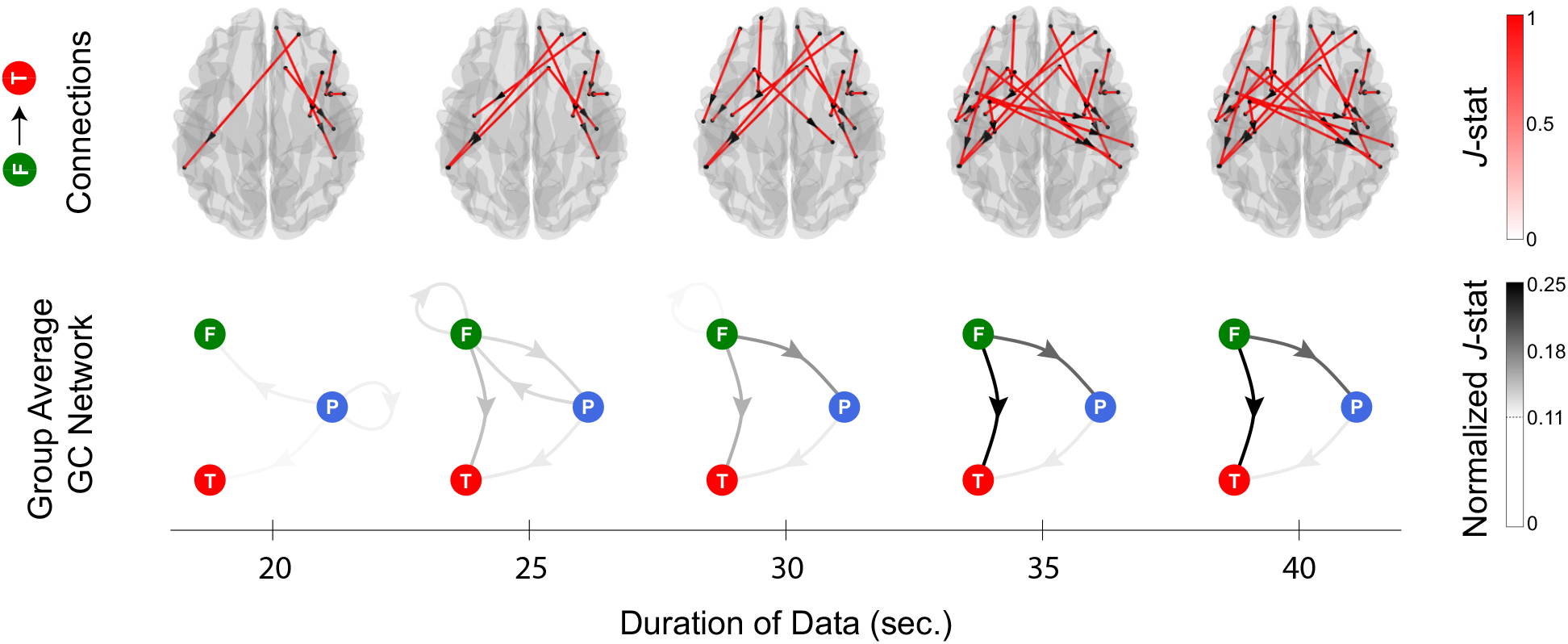
Evaluating the effect of trial duration on the NLGC performance. The group average GC links from frontal to temporal areas for younger participants during tone processing are overlaid on the dorsal brain plot in the top tow. The corresponding directed graphs indicating the normalized *J*-statistics of the links between frontal, temporal, and parietal areas are shown in the bottom row. Columns correspond to different choices of *T* corresponding to the first 20, 25, 30, 35 and 40 s of the data. While for smaller values of *T*, several links are missing, by increasing *T* beyond 30 s the detected networks stabilize and converge.

#### 4.8.4. Statistical Testing

We used generalized linear mixed effect models (GLMM) to analyze the effects of age, condition, connectivity type and hemisphere on the GC link counts for each frequency band. The statistical analysis was conducted via R version 4.0.5 (R core Team 2021) using glmmTMB (Brooks et al., 2017) with zero-inflated generalized Poisson distributions to model the link counts. Based on a full model accounting for all the variables, the best fit model was selected by stepwise elimination, implemented in buildglmmTMB Voeten (2021) based on the likelihood ratio test (LRT). Model assumptions for dispersion, heteroskedasticity and zero-inflation were examined and verified using the DHARMa package (Hartig, 2021). The post-hoc differences among the levels of the effects were tested using pairwise comparisons based on estimated marginal means, with Holm corrections using the package emmeans Lenth (2021). The summary of the statistical models is given in Appendix C.

## 5. Acknowledgments

This work was supported in part by the National Science Foundation Awards No. OISE2020624, SMA1734892 and CCF1552946 and the National Institutes of Health Awards. No. R01-DC019394, R01-DC014085, P01-AG055365, and R21-AG068802.

## Appendix A. Parameter Estimation

This appendix provides the details and derivations of the EM algorithm used in NLGC as well as the VAR fitting used by the two-stage approaches. The EM algorithm is derived in Appendix A.1. In Appendix A.2, we present the filtering and smoothing procedures to obtain the conditional distribution *p*(**x**_1:*T*_|**y**_1:*T*_; ***θ***), followed by the VAR fitting procedure used in two-stage approaches that are derived in Appendix A.3.

### Appendix A.1. EM Algorithm

In this section, we derive the E- and M-steps used in the network parameter estimation module of NLGC.

#### E-step

We start from the joint distribution of 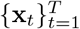 and 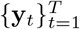. From the Bayes’ rule we have

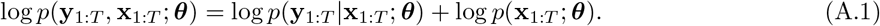

The conditional distribution can be directly written from observation model in Eq. (1) as

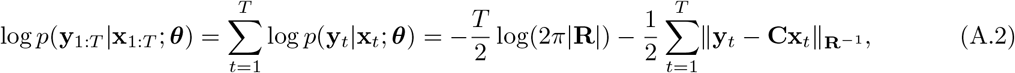

where ||**a**||_**B**_ := **a^⊤^Ba** is utilized for notational convenience.

Using the fact that 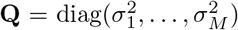 along with the source dynamic model in Eq. (2), one can write down

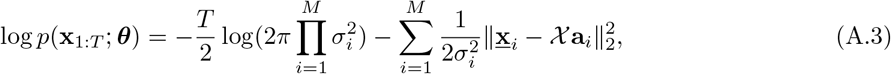

where 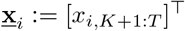, **a**_*i*_ = [[**A**_*k*_]_*i,j*_, ∀*k,j*]^⊤^, and

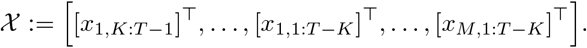

Now, substituting Eqs. (A.2) and (A.3) into Eq. (A.1) along with taking the expectation yields

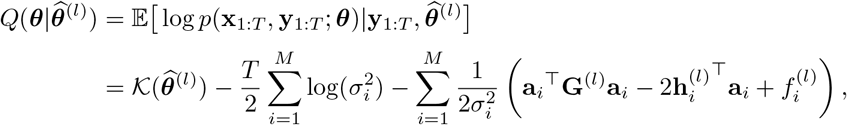

where 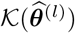 represents the constant terms with respect to ***θ***

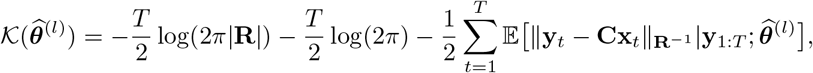

and

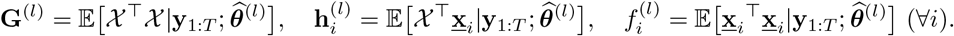

It is noteworthy to mention that the variables **G**^(*l*)^, 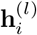, and 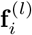 can be written as a function of first- and second-order moments of the conditional density 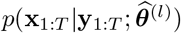. It can be shown that the conditional density 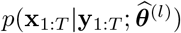 is Gaussian due to underlying Gaussian assumptions on **w**_*t*_ and **n**_*t*_. Thus, the mean and covariance matrices can be efficiently computed via the Fixed Interval Smoothing (FIS) algorithm (Anderson and Moore, 2005). The details are presented in the next subsection.

#### M-step

To avoid ill-posedness caused by the low-dimensional MEG measurements, we leverage the sparse connectivity feature of cortical sources and add a regularization term in the M-step as follows:

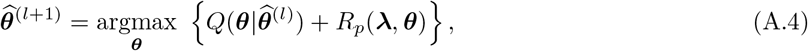

where 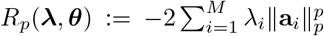 is the regularization function and **λ** = [λ_1_,…,λ_*M*_]^⊤^ ∈ ℝ^*M*^ is the regularization coefficients vector. The closed-form solution for *p* = 2 can be obtained as

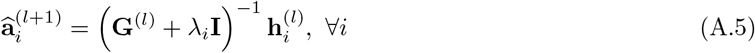

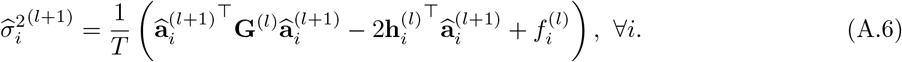

To enforce sparsity, we use *p* = 1. However, the closed-form solution does not exist. We use the well-known *Fast Adaptive Shrinkage/Thresholding Algorithm* (FASTA) to find the ℓ_1_-norm regularized solution to Eq. (A.4) (Goldstein et al., 2014).

The EM procedure for the full model is summarized in Algorithm 2. It is noteworthy that in order to find the VAR model parameters for the full and reduced models, one requires to run the algorithm for a total of *M*(*M* – 1) + 1 times, i.e., 1 full model where we consider all interactions between the sources and *M*(*M* – 1) reduced models corresponding to all possible links in the set 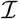. Thus, it is crucial to have computationally efficient solutions to carry out the computations in the E-step. Before presenting the FIS procedure used for this purpose, some remarks regarding the initialization of the EM algorithm, estimating the reduced models, and choosing the regularization parameters **λ** are in order:

#### Remark 1.

(**Initialization**) Due to the biconvex nature of the problem in Eq. (A.4), the problem may have several saddle points. As a result, choosing a proper initial point for the EM algorithm is crucial and helps the algorithm to converge faster as well. We first obtain the minimum norm source estimates as follows

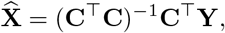

where 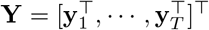 is the MEG measurement matrix and 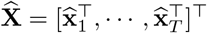 is the source estimates matrix. Given the source estimates, we initialize all coefficients 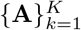 with zero and variances matching the average power of each source, i.e., 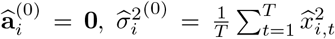, ∀*i*. In this way, the algorithm is initialized with an unbiased solution (Gorodnitsky et al., 1995).

#### Remark 2.

(**Reduced Models**) Algorithm 2 represents the full model parameter estimation. With some minor modification, one can find the reduced model estimation in a similar way. Let us assume we want to estimate the reduced model parameters corresponding to the link 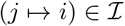. We can use Algorithm 2 by enforcing **a**_*i,j,k*_ = 0, ∀*k* at the M-step in each iteration. The output of the Algorithm 2 in this case is the estimated parameters for the reduced model corresponding to the link (*j* ↦ *i*).

**Algorithm 2.**
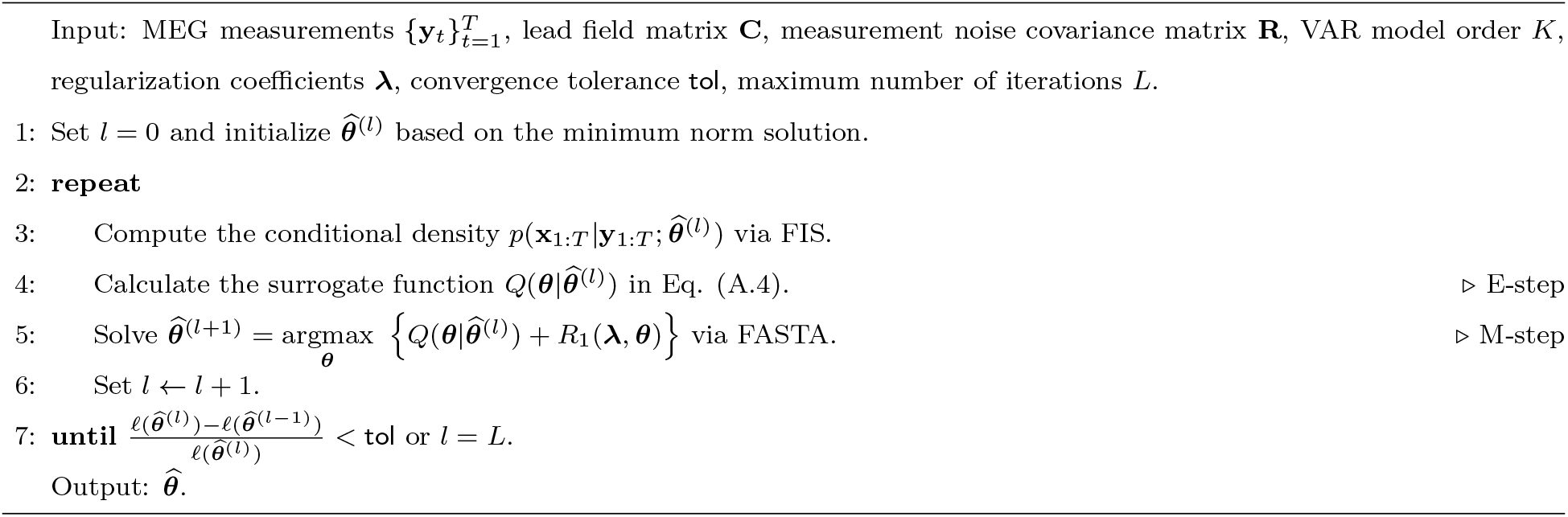
EM-based Parameter Estimation

#### Remark 3.

(**Regularization Parameters)** To obtain the regularization parameters **λ**, we utilize the standard *K*-fold cross-validation. To save the computational complexity and to speed up the tuning process, we assume **λ** = λ**1** where **1** is the all-one vector. As for the cross-validation metric, we use the estimation stability criterion presented in (Lim and Yu, 2016). Given a set of candidates for λ, this criterion constructs estimated versions of the MEG measurements based on the underlying parameters of the VAR model and returns the model with the lowest variance across the folds. In this way, the chosen λ gives a stable solution across the folds. Moreover, once the optimal regularization parameter λ is chosen for the full model, we use the same regularization parameter for all the subsequent reduced models (Das and Babadi, 2021). This way, the cross-validation only needs to be carried out for the full model.

### Appendix A.2. Fixed Interval Smoothing

As mentioned earlier, under Gaussian assumptions on **n**_*t*_ and **w**_*t*_, the conditional density of *p*(**x**_1:*T*_|**y**_1:*T*_; ***θ***) is also Gaussian (Anderson and Moore, 2005). As a result, we just need to find the conditional mean and covariance matrix of the random vector **x**_1:*T*_ given **y**_1:*T*_ and ***θ***.

Using the Kalman filter, we can compute the filtered densities *p*(**x**_*t*_|**y**_1:*t*_; ***θ***) for *t* = 1, 2,…, *T*. Using the filtered densities, the FIS procedure allows us to also find *p*(**x**_*t*_|**y**_1:*T*_; ***θ***) for *t* = 1, 2,…, *T*. To this end, we first perform state augmentation to transform VAR(*K*) models to equivalent VAR(1) models. The augmented state vector is defined as 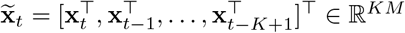. The VAR(*K*) model in Eq. (2) can thus be rewritten as a VAR(1) model given by:

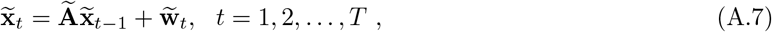

where

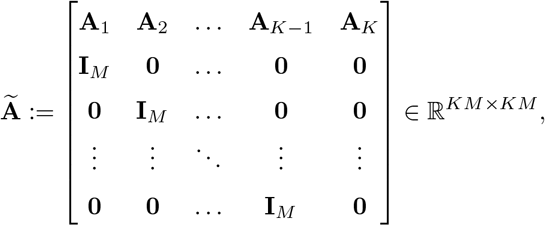

and 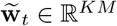 is the augmented state noise vector with covariance matrix 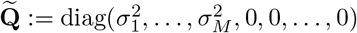. Similarly, we can modify the measurement model in Eq. (1) as follows

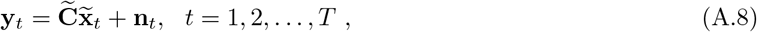

with 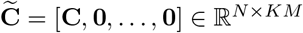.

Let us define the conditional mean, covariance, and cross-variance of the sources as 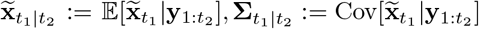, and 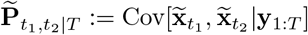, respectively, for two given time-points 1 ≤ *t*_1_, *t*_2_ ≤ *T*. Assuming that matrices 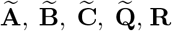, and 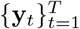 are given, we can utilize the *Kalman filter* to obtain 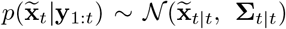, *t* = 1,…,*T*. Next, we use FIS to also find 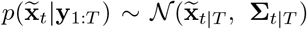, *t* = 1,…,*T*.

According to (Jong and Mackinnon, 1988), for the the conditional cross-covariance, we have the following recursive relationship:

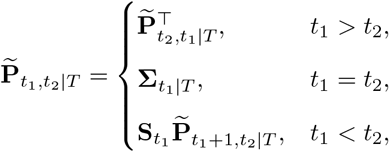

where 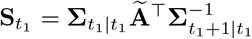.

Finally, to extract the first- and second-order moments of the sources from the augmented model, we define 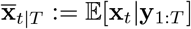 and **P**_*t*_1_*t*_2_|*T*_ := Cov[**x**_*t*_1__, **x**_*t*_2__ |**y**_1:*T*_]. From the definition of the augmented model, we have

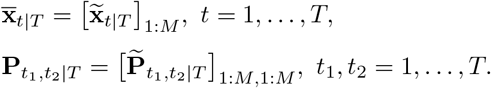

Algorithm 3 summarizes the overall procedure for finding the smoothed means and covariance matrices. A costly computational step in Algorithm 3 is the inversion of **∑**_*t*+1|*t*_ ∈ ℝ^*KM*×*KM*^ that needs to be performed in each iteration. In order to mitigate this source of computational complexity, we use the steady-state filtering approach of (Pirondini et al., 2018). Let us define the steady-state covariance matrices **Σ**^(+)^ and **Σ**^(-)^ as follows

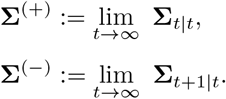

**Algorithm 3.**
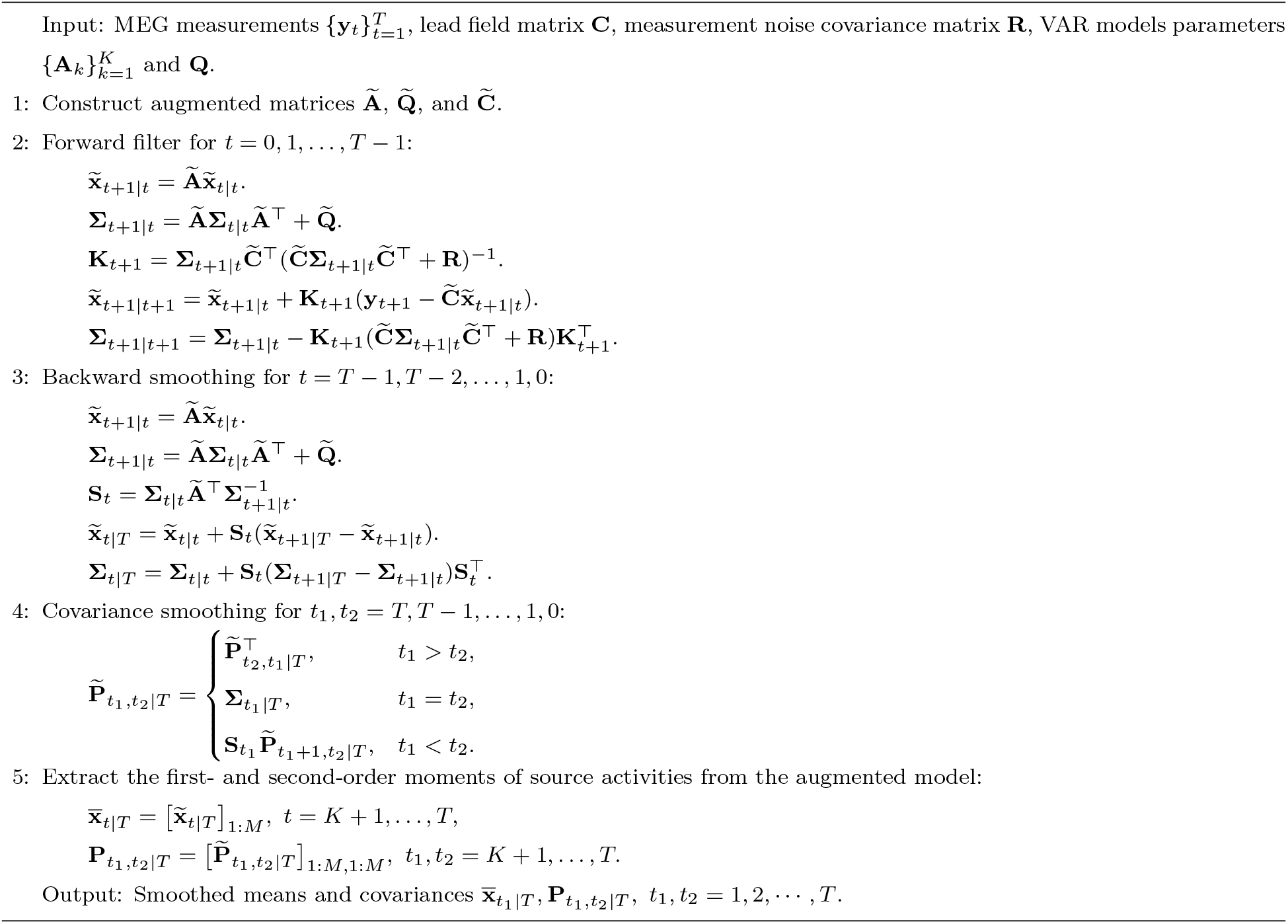
Fixed Interval Smoothing

Replacing these steady-state values into the forward filter yields

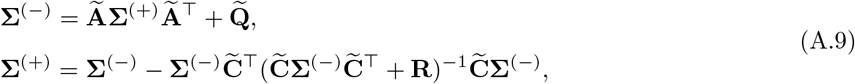

which is known as the discrete-time algebraic Riccati (DARE) equation with respect to **Σ**^(+)^. The DARE equation can be solved efficiently using the MacFarlane-Potter-Fath eigen-structure method (Malik et al., 2011). Solving the Riccati equation gives the steady-state covariance matrices and from there, we can compute the Kalman gain (**K**_*t*_) and smoothing gain (**S**_*t*_) independent of *t*:

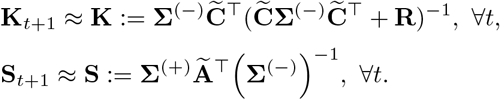

As a result, only two matrix inversions are required at the beginning of the FIS, thereby providing significant computational savings.

### Appendix A.3. VAR Model Fitting in the Two-Stage Methods

In the two-stage approaches, the source estimates are first computed using a source localization procedure, followed by VAR model fitting. Let us denote the source estimates by 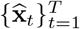. The VAR(*K*) model fitting can be performed in various ways, among which maximum likelihood estimation is a popular method (Haykin, 2013). To this end, one needs to compute 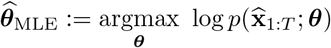, where

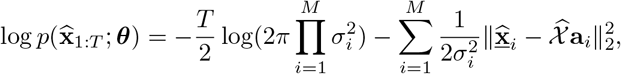

with 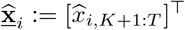, and 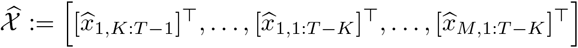. Setting the derivative of the log-likelihood with respect to the parameters to zero gives the following closed-form solution

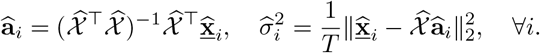

Similar to NLGC, we can enforce sparsity by considering an ℓ_1_-norm regularized maximum likelihood problem. To this end, we need to find 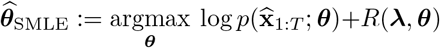, where 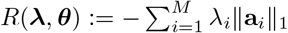 is the ℓ_1_-norm penalty and 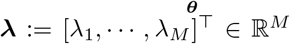 is the regularization vector. As mentioned in Appendix A.1, this problem does not have a closed-form solution. However, we can use iterative methods such as FASTA (Goldstein et al., 2014) or *Iteratively Re-weighted Least S’quares* (IRLS) (Ba et al., 2014) to obtain the ℓ_1_-norm regularized estimates. The regularization parameters **λ** can be tuned using standard cross-validation techniques, as mentioned in Appendix A.1.

## Appendix B. Proof of Theorem 1

The proof of Theorem 1 follows that of the main theorem in (Sheikhattar et al., 2018). First, we define the following notations for a given log-likelihood function ℓ(***θ***) with parameter ***θ***:

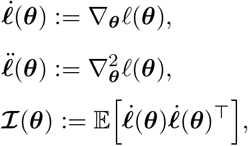

where 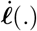 denotes the gradient vector of the likelihood with respect to ***θ***, also referred to as the score statistics, 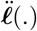 denotes the Hessian matrix of the log-likelihood, and 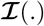 is the Fisher information matrix. We define the de-biased deviance difference between the true value of ***θ*** and its estimate 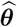 as (Sheikhattar et al., 2018):

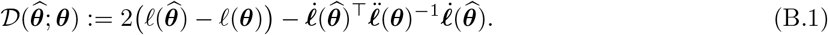

Starting from the definition of the log-likelihood function, we can decompose ℓ(***θ***) as

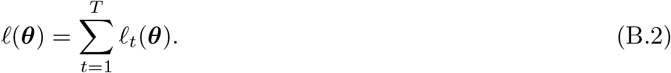

where ℓ_*t*_(***θ***) = log *p*(**y**_*t*_|**y**_1:*t*–1_; ***θ***) for *t* = 2,⋯, *T* with the convention ℓ(***θ***) = log *p*(**y**_1_; ***θ***). Using the second-order Taylor expansion of ℓ(***θ***) around 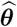 along with the intermediate value theorem, we have

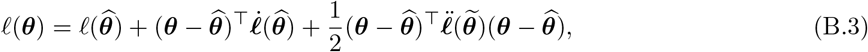

where 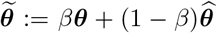 for some *β* ∈ (0,1) such that 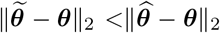. Substituting ℓ(***θ***) from Eq. (B.3) into Eq. (B.1) gives

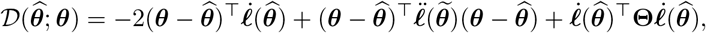

where 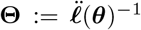. Using an auxiliary vector 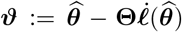 and after rearrangement, the de-biased deviance can be rewritten as

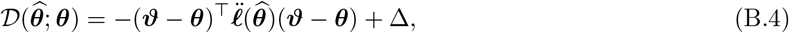

with

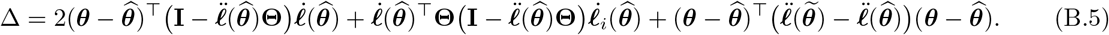

Employing the consistency of the estimation, i.e., 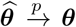 and the Lipschitz property of the second-order derivative of the Gaussian log-likelihood function, one can show that the term Δ asymptotically goes to zero as *T* → ∞ with a rate of 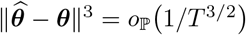 (van de Geer et al., 2014; Sheikhattar et al., 2018).

Let us now consider the link 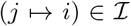. In what follows, we prove the first and second assertions of the theorem regarding the null and alternative hypotheses separately.

### Null Hypothesis

The Taylor expansion of the score statistics can be expressed as

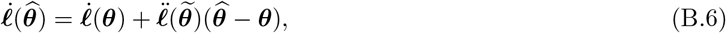

where 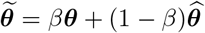 for some *β* ∈ (0,1). Combining the Taylor expansion in Eq. (B.6) along with the definition 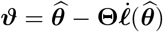, we have

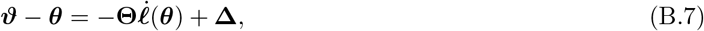

with 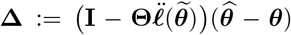. Following the same argument for Δ in Eq. (B.5), one can show that 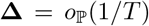 is asymptotically negligible as *T* → ∞ (van de Geer et al., 2014). In order to obtain the asymptotics of the score statistic and the Hessian matrix of the log-likelihood function ℓ(***θ***), the conventional law of large numbers (LLN) and the central limit theorem (CLT) can be used, since the process realizations in the log-likelihood decomposition of Eq. (B.2) (**y**_*t*_|**y**_1:*t*–1_, ∀*t* > 1) are independent across time. This is due to the fact that the noise processes **w**_*t*_ and **n**_*t*_ in our generative model are i.i.d. Gaussian noise sequences and are independent of each other (Anderson and Moore, 2005).

Using the LLN for the Hessian matrix of ℓ(·) yields

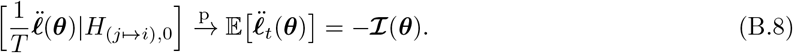

Moreover, the CLT for the score statistics gives

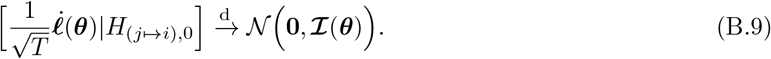

Using Slutsky’s theorem along with Eqs. (B.6), (B.8), and (B.9), asymptotic normality of ***ϑ*** under the null hypothesis can be obtained as

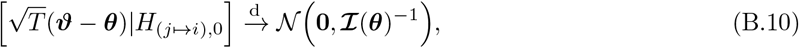

as *T* ↦ ∞. Following the definition of the deviance in Eq. (B.4) along with Eq. (B.8), we have

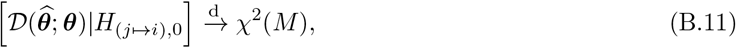

as *T* ↦ ∞, where *M* is the dimension of the parameter ***θ***. Following the results in (Wald, 1943) and (Wilks, 1938) along with the fact that 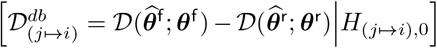, it can be shown that the de-biased deviance difference converges to a central *χ*^2^ distribution with *M*^d^ degrees of freedom

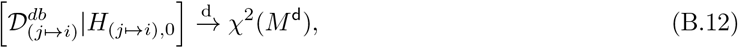

where *M*^d^ = *M*^f^ – *M*^r^ is the difference between dimensions of the two nested models. This proves the first assertion of Theorem 1.

### Alternative Hypothesis

Following the development in (Davidson and Lever, 1970), we define a non-decreasing sequence 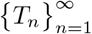 such that lim_*n*↦∞_ *T_n_* = *T*. Instead of defining a fixed alternative against the null hypothesis *H*_(*j*↦*i*),0_ : ***θ*** = (***θ***_0_, **0**), we instead define a sequence of local alternatives

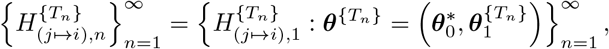

where 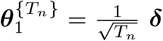 is an unspecified sub-vector excluded from the reduced model with dimension *M*^d^ = *M*^f^ – *M*^r^ and ***δ*** is a constant vector. According to (Davidson and Lever, 1970), we test for the departure of the sequence of local alternatives from the null hypothesis at the true parameter 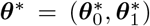 with 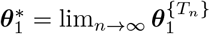.

For notational convenience, we hereafter drop the subscript *n* in *T_n_*, noting that the equations involving limits of *T* denote sequential limits. Defining the de-biased vector 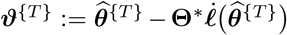 corresponding to the local alternative 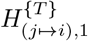 with 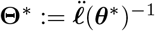 and utilizing the following expansions

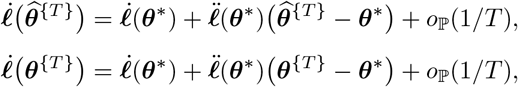

we have

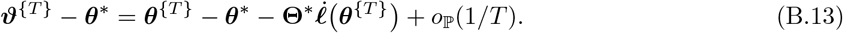

Using LLN and CLT similar to the case of the null hypothesis, we conclude

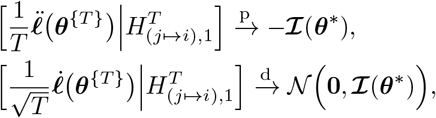

and the asymptotic normality of ***ϑ*** follows as

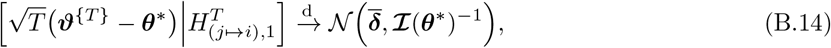

where 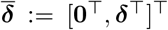 is the asymptotic mean. It is noteworthy that the non-zero asymptotic mean is obtained from the *Pitman drift* rate where the sequence of true local parameters ***θ***^{*T*}^ tends to its limit ***θ***\* at a rate 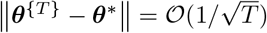 (Davidson and MacKinnon, 1987).

Next, using an extension of *Cochrans theorem* to non-central chi-square distribution (Tan, 1977) and using the asymptotic normality of ***ϑ***^{*T*}^ in Eq. (B.14), it follows that under the sequence of local alternatives 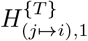, the de-biased deviance difference of the two nested full and reduced models converges to a non-central chi-squared distribution as *T* ↦ ∞:

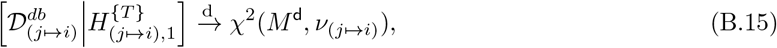

where *M*^d^ is the difference between the dimensions of the two nested models and *ν*_(*j*↦*i*)_ presents the noncentrality parameter. To identify the non-centrality parameter, let us consider the block decomposition of 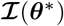 corresponding to 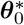 and 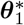 as

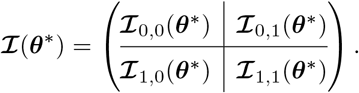

**Figure B.1:**
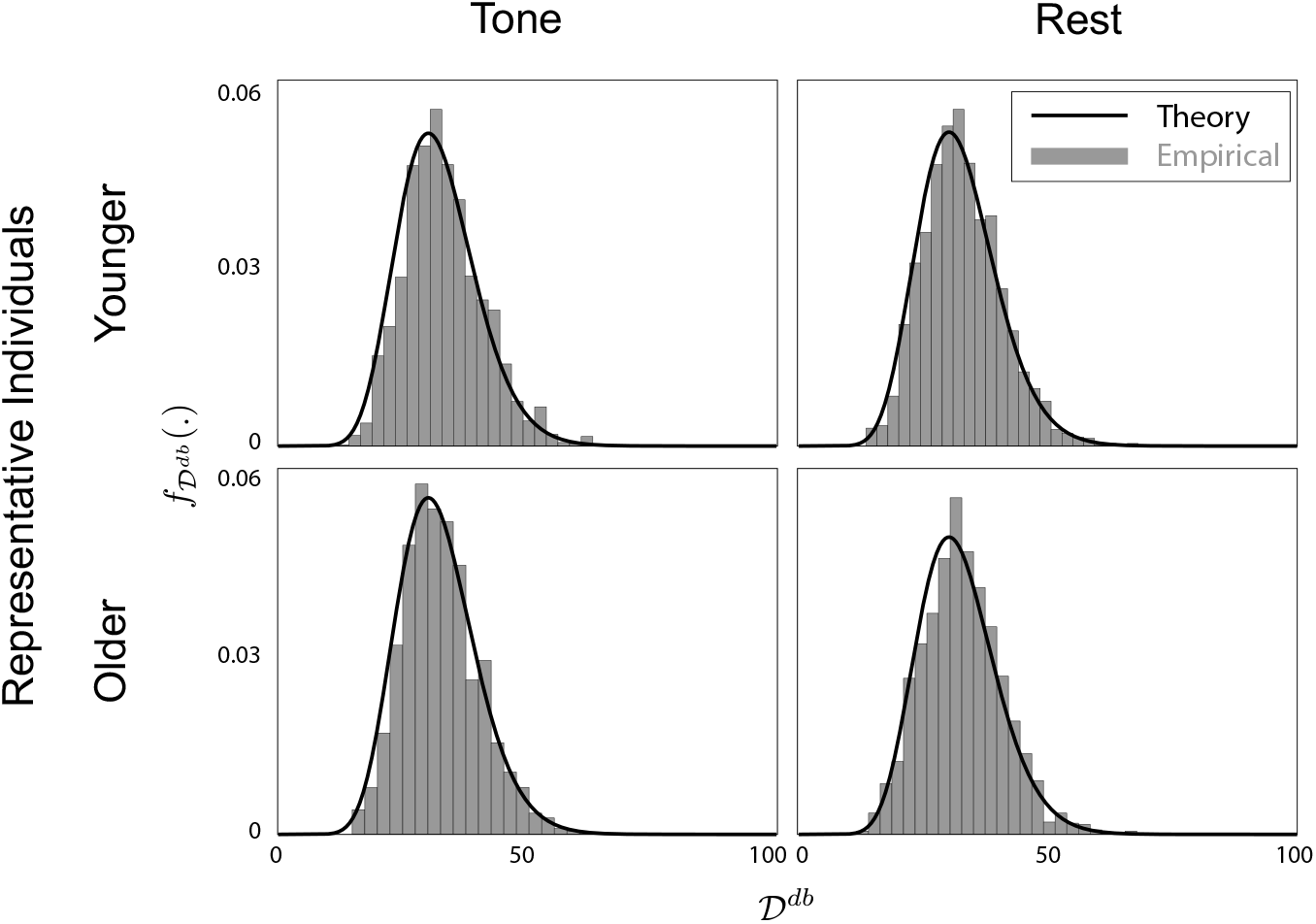
Histograms of the debiased deviance differences corresponding to non-GC links for younger and older representative subjects in tone and rest conditions from Section 2.4. The histograms closely match the prediction of Theorem 1.

Then, 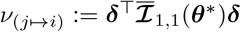 with 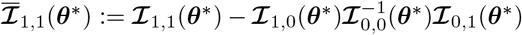. This proves the second assertion of the theorem.

Finally, to test whether the theoretical prediction of Theorem 1 regarding the null distribution is valid for our analysis of experimental MEG data, we chose 4 representative trials (one older and one younger participant in each condition) and plotted the histogram of the debiased deviance differences of all the tested GC links that were not significant. According to Theorem 1, the debiased deviance differences of such non-GC links should follow a chi-square distribution with degree of freedom 2 × 4^2^ = 32 (*r* = 4 eigenmodes and VAR(2) model). Fig. B.1 shows the corresponding chi-square density and the empirical histograms. As it can be seen, the empirical histograms closely match the theoretical chi-square density.

## Appendix C. Mixed-Effects Model

Full models for the mixed effect models included interactions among the fixed effects of age, condition, connectivity type and hemisphere, and random slopes and intercepts for within-subject factors of condition, connectivity type and hemisphere per subject. Summary tables for each frequency band are given in Table C.1.

**Table C.1:** Statistical model summary table corresponding to Section 2.4.

| Parameter | Delta+Theta Band | Beta Band |
| --- | --- | --- |
| Count model: (Intercept) | 3.06(0.07)*** | 2.24(0.10)*** |
| Count model: connectivityF->P | -0.84(0.11)*** | -0.10(0.14) |
| Count model: connectivityF->T | -1.29(0.13)*** | 0.29(0.12)* |
| Count model: connectivityP->F | 0.13(0.08) | 0.96(0.11)*** |
| Count model: connectivityP->P | -0.79(0.12)*** | 1.03(0.11)*** |
| Count model: connectivityP->T | -0.84(0.11)*** | 0.86(0.11)*** |
| Count model: connectivityT->F | -0.29(0.09)** | 0.66(0.12)*** |
| Count model: connectivityT->P | -1.10(0.12)*** | -0.02(0.13) |
| Count model: connectivityT->T | -0.97(0.12)*** | -0.13(0.14) |
| Count model: AgeOlder | -0.16(0.11) | -0.05(0.16) |
| Count model: Conditiontone | -0.93(0.12)*** | 0.96(0.11)*** |
| Count model: hemi2inter | -0.01(0.04) |  |
| Count model: connectivityF->P:AgeOlder | -0.18(0.18) | -0.10(0.23) |
| Count model: connectivityF->T:AgeOlder | 0.25(0.21) | -0.30(0.22) |
| Count model: connectivityP->F:AgeOlder | 0.05(0.13) | -0.07(0.19) |
| Count model: connectivityP->P:AgeOlder | 0.26(0.17) | -0.29(0.19) |
| Count model: connectivityP->T:AgeOlder | -0.46(0.21)* | 0.24(0.18) |
| Count model: connectivityT->F:AgeOlder | 0.12(0.15) | -0.42(0.20)* |
| Count model: connectivityT->P:AgeOlder | 0.26(0.19) | -0.25(0.23) |
| Count model: connectivityT->T:AgeOlder | 0.14(0.18) | -0.03(0.23) |
| Count model: connectivityF->P:Conditiontone | 1.86(0.16)*** | -0.99(0.19)*** |
| Count model: connectivityF->T:Conditiontone | 2.61(0.17)*** | -0.88(0.16)*** |
| Count model: connectivityP->F:Conditiontone | -0.07(0.16) | -1.31(0.15)*** |
| Count model: connectivityP->P:Conditiontone | 1.39(0.17)*** | -1.65(0.15)*** |
| Count model: connectivityP->T:Conditiontone | 1.47(0.16)*** | -1.60(0.16)*** |
| Count model: connectivityT->F:Conditiontone | -0.07(0.17) | -1.07(0.15)*** |
| Count model: connectivityT->P:Conditiontone | 1.13(0.18)*** | -0.82(0.17)*** |
| Count model: connectivityT->T:Conditiontone | 0.91(0.19)*** | -0.82(0.18)*** |
| Count model: AgeOlder:Conditiontone | -0.50(0.22)* | -0.57(0.19)** |
| Count model: Conditiontone:hemi2inter | -0.32(0.06)*** |  |
| Count model: connectivityF->P:AgeOlder:Conditiontone | 0.51(0.29) | 1.57(0.29)*** |
| Count model: connectivityF->T:AgeOlder:Conditiontone | 0.30(0.30) | 0.46(0.33) |
| Count model: connectivityP->F:AgeOlder:Conditiontone | 0.72(0.28)* | 0.22(0.26) |
| Count model: connectivityP->P:AgeOlder:Conditiontone | 0.64(0.28)* | 0.90(0.26)*** |
| Count model: connectivityP->T:AgeOlder:Conditiontone | 1.20(0.31)*** | 0.43(0.26) |
| Count model: connectivityT->F:AgeOlder:Conditiontone | 1.02(0.29)*** | 0.77(0.26)** |
| Count model: connectivityT->P:AgeOlder:Conditiontone | 0.40(0.32) | 0.26(0.32) |
| Count model: connectivityT->T:AgeOlder:Conditiontone | 0.67(0.32)* | 1.03(0.29)*** |
| Zero model: (Intercept) | -3.49(0.22)*** | -3.31(0.18)*** |
| AIC | 10122.64 | 10803.40 |
| Log Likelihood | -5020.32 | -5362.70 |
| Num. obs. | 1584 | 1584 |
| Num. groups: MEG_ID | 22 | 22 |
| Var (count model): MEG_ID (Intercept) | 0.01 | 0.01 |

\*\*\* $p < 0.001$ ; \*\* $p < 0.01$ ; \* $p < 0.05$

